# Reciprocal links between methionine metabolism, DNA repair and therapy resistance in glioblastoma

**DOI:** 10.1101/2024.11.20.624542

**Authors:** Navyateja Korimerla, Baharan Meghdadi, Isra Haq, Kari Wilder-Romans, Jie Xu, Nicole Becker, Ziqing Zhu, Peter Kalev, Nathan Qi, Charles Evans, Maureen Kachman, Zitong Zhao, Angelica Lin, Andrew J Scott, Alexandra O’Brien, Ayesha Kothari, Peter Sajjakulnukit, Li Zhang, Sravya Palavalasa, Erik R Peterson, Marc L Hyer, Katya Marjon, Taryn Sleger, Meredith A Morgan, Costas A Lyssiotis, Everett M. Stone, Sean P. Ferris, Theodore S Lawrence, Deepak Nagrath, Weihua Zhou, Daniel R Wahl

**Author notes:** Corresponding author, To whom correspondence should be addressed: Daniel R. Wahl, MD, PhD Department of Radiation Oncology University of Michigan, UH B2 C490, Ann Arbor, MI 48109.

## Abstract

Glioblastoma (GBM) is uniformly lethal due to profound treatment resistance. Altered cellular metabolism is a key mediator of GBM treatment resistance. Uptake of the essential sulfur-containing amino acid methionine is drastically elevated in GBMs compared to normal cells, however, it is not known how this methionine is utilized or whether it relates to GBM treatment resistance.

Here, we find that radiation acutely increases the levels of methionine-related metabolites in a variety of treatment-resistant GBM models. Stable isotope tracing studies further revealed that radiation acutely activates methionine to S-adenosyl methionine (SAM) conversion through an active signaling event mediated by the kinases of the DNA damage response. *In vivo* tumor SAM synthesis increases after radiation, while normal brain SAM production remains unchanged, indicating a tumor- specific metabolic alteration to radiation. Pharmacological and dietary strategies to block methionine to SAM conversion slowed DNA damage response and increased cell death following radiation in vitro. Mechanistically, these effects are due to depletion of DNA repair proteins and are reversed by SAM supplementation. These effects are selective to GBMs lacking the methionine salvage enzyme methylthioadenosine phosphorylase. Pharmacological inhibition of SAM synthesis hindered tumor growth in flank and orthotopic *in vivo* GBM models when combined with radiation. By contrast, methionine depletion does not reduce tumor SAM levels and fails to radiosensitize intracranial models, indicating depleting SAM, as opposed to simply lowering methionine, is critical for hindering tumor growth in intracranial models of GBM.

These results highlight a new signaling link between DNA damage and SAM synthesis and define the metabolic fates of methionine in GBM *in vivo*. Inhibiting radiation-induced SAM synthesis slows DNA repair and augments radiation efficacy in GBM. Using MAT2A inhibitors to deplete SAM may selectively overcome treatment resistance in GBMs with defective methionine salvage while sparing normal brain.

## INTRODUCTION

Glioblastoma is a lethal form of brain cancer driven by numerous metabolic abnormalities^1–4^. Most GBM patients are treated with radiotherapy (RT) and chemotherapy, but tumors inevitably recur.^5^ Our group and others have found that the metabolic abnormalities that fuel GBM also drive this treatment resistance^6–8^, yet we lack a full understanding about how to disrupt these links to improve patient outcomes.^9^

Methionine metabolism is dramatically altered in GBM^10^. Methionine is a sulfur-containing essential amino acid that drives various cellular functions such as methylation, polyamine synthesis, redox balance, and nucleotide synthesis. GBMs have elevated methionine consumption compared to cortex, which makes methionine-based imaging strategies useful for gliomas^11–14^. These imaging modalities allow the measurement of methionine *uptake*, but we have no knowledge regarding which downstream methionine-*consuming* pathways are utilized in GBM, how these differ from those utilized by the normal cortex, if these are influenced by treatments such as RT, and if they mediate glioma treatment resistance.

S-adenosyl methionine (SAM), the universal methyl donor, is an important methionine-derived metabolite that drives a variety of transmethylation reactions. While SAM-driven methylation reactions can promote DNA repair, the complete mechanistic links between DNA damage, methionine metabolism and DNA repair are unknown^15–17^. Thus, while methionine depletion has been exploited therapeutically to inhibit tumor growth on its own and in combination with genotoxic therapy^18,19^, we do not have an understanding of how best to target this biology to improve therapeutic responses in GBM.

In the present work, we define links between methionine metabolism, SAM formation, DNA repair, and RT resistance in GBM. We find that RT acutely increases the abundance of methionine-related metabolites in RT-resistant GBM models. Using stable isotope tracing and molecular biology techniques, we identify that RT acutely increases SAM synthesis in GBM *in vitro* and *in vivo* through a DNA damage response-mediated signaling mechanism. Interfering with methionine metabolism by depleting SAM delays DNA repair and increases the sensitivity of GBM models to RT. These effects are most pronounced in GBM models that cannot salvage SAM. These findings indicate that inhibiting SAM synthesis (which is being clinically investigated in other cancers) may be a promising strategy to overcome therapy resistance in GBM.

## RESULTS

### RT acutely alters methionine metabolism in glioblastoma

Gliomas, including the aggressive brain cancer glioblastoma (GBM), take up increased methionine compared to the normal cortex, however it is not known how this methionine is utilized or whether it relates to the profound treatment resistance of these cancers.^11,20,21^ To begin to understand these links, we quantified the levels of methionine-related metabolites in normal mouse brain and xenograft GBM tumor tissue. We utilized a patient derived GFP-expressing classical transcriptional subtype and treatment resistant GBM38 model, to form orthotopic tumors in mice.^22^ Using fluorescent guided microdissection, we isolated the GFP-expressing tumor tissue from the contralateral brain. Methionine metabolism can drive a variety of downstream pathways like transmethylation, antioxidant synthesis, methionine salvage and polyamine synthesis (Fig. 1A). Methionine was elevated 5-fold in orthotopic brain tumors compared to the contralateral brain (Fig. 1B). Several methionine-related metabolites involved in these pathways including S-adenosyl methionine (SAM), cystathionine, glutathione and spermidine are also elevated in GBMs when compared to the contralateral brain (Fig. 1B). We confirmed these findings using another orthotopic patient derived mesenchymal transcriptional subtype and treatment resistant HF2303 GBM tumor model where similar methionine related metabolites involved in transmethylation, antioxidant synthesis and polyamine synthesis exhibited a comparable trend of elevated levels in GBM compared to contralateral brain (Fig. S1A).^23^

**Figure 1.**
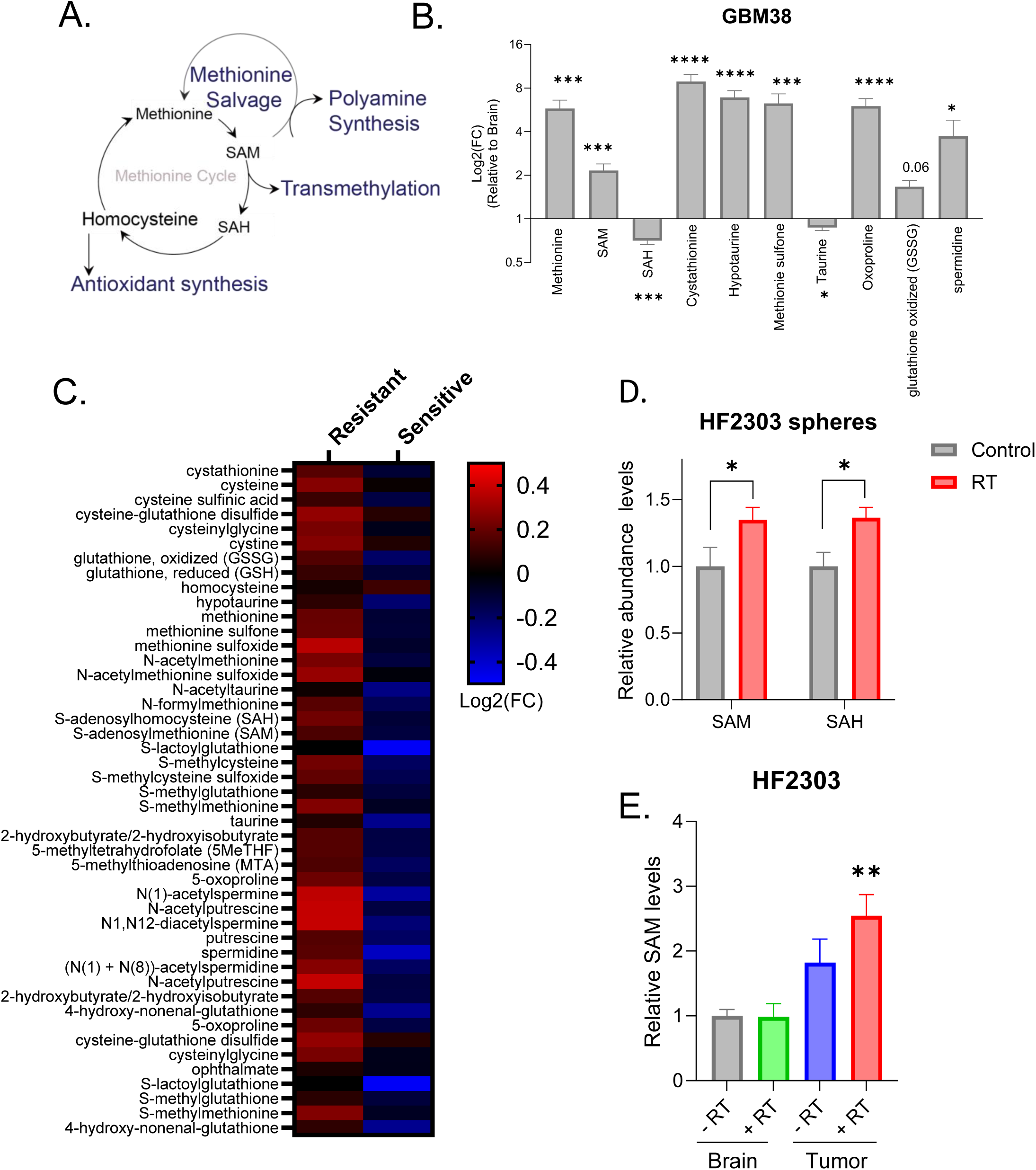
RT acutely alters GBM methionine metabolism. A). Schema showing the metabolic pathways associated with methionine metabolism. B). Untreated intracranial GBM38 xenografts and the contralateral mouse brain were collected, and methionine metabolites were measured using LC/MS. Data represented as mean±SEM for N=6 biological replicates compared with the control (Brain). C). Two radiation therapy-resistant (U87 MG and A172) and two radiation therapy-sensitive (KS- 1 and U118 MG) GBM cell lines were irradiated with 8 Gy, and harvested 2 h after RT. The samples were analyzed by targeted LC-MS/MS (4 biologic replicates per cell line) to determine fold-change values for each metabolite based on unirradiated matched cell line controls. Heatmap represents the average fold change for RT- resistant (left) and RT-sensitive cell lines (right). D). Patient-derived HF2303 neurospheres were irradiated with 8 Gy and total SAM and SAH levels were analyzed using MS, 1 hour after RT. Data represented as mean±SEM for N=8 biological replicates. E). HF2303 PDX-carrying mice were treated with or without cranial RT (2*2 Gy), and the tumor and normal brain were harvested 4-hour post the last RT dose and analyzed using LC-MS. Data represented as mean±SEM for N=9-12 biological replicates. P values were obtained in comparison with the control (Brain-RT). P of Tumor+RT compared to Tumor-RT is 0.0758. A-E). *P ≤ 0.05, **P ≤ 0.01, ***P ≤ 0.001 and ****P ≤ 0.0001.

We hypothesized these changes in methionine metabolism in GBM tumors can influence their response to RT, thereby conferring therapy resistance. To address this question, we interrogated the abundances of methionine-related metabolites in untreated and irradiated GBM cells in an existing data set.^6^ We found that RT increases the abundance of methionine and several methionine-related metabolites involved in transmethylation, antioxidant synthesis, methionine salvage and polyamine synthesis in two different RT-resistant immortalized GBM cell lines, A172 and U87(Fig. 1C, left column). Heatmap represents the average fold change for RT-resistant (A172 and U87) and RT-sensitive cell lines (KS1 and U118MG) for each metabolite based on unirradiated matched cell line controls. However, the RT-induced increase in metabolite abundances was not observed in RT-sensitive cell lines, KS1 and U118MG (Fig.1C, right column). These findings further supported our hypothesis that GBMs not only have elevated methionine and methionine related metabolites, but these metabolites are differentially altered in RT resistant vs sensitive cell lines, indicating that methionine related metabolites might influence the response to RT.

We then sought to understand if RT altered methionine metabolism in GBM models that were more relevant to human tumors, including neurospheres and intracranial tumors.^24^ We focused on SAM, the immediate downstream product of methionine, and SAH, which is formed when SAM donates a methyl group to a transmethylation reaction (Fig. 1A). We irradiated patient derived HF2303 neurospheres and found that RT acutely increases both SAM and SAH abundance (Fig. 1D). RT also acutely increased SAM levels in intracranial HF2303 tumors but did not alter SAM levels in the contralateral cortex (Fig. 1E). We observed similar and significant changes in SAH (Fig. S1B). Methionine and 5’-Methylthioadenosine (MTA) levels are higher in tumors compared to the contralateral cortex but are unaffected by RT (Fig. S1C and Fig. S1D). These results were recapitulated in mice bearing GBM6 PDXs (a classical subtype treatment resistant PDX from Mayo Clinic GBM resource), where SAM levels were higher in tumor than cortex, and increased further following RT in tumor, but not cortex (Fig. S1E). We observed similar, but non-significant changes in GBM tumors grown in mouse flanks (Fig. S1F). Thus, we conclude that RT therapy rapidly alters methionine metabolism, specifically acutely elevating SAM and SAH levels in glioblastoma models *in vitro* and *in vivo*.

### RT acutely increases SAM synthesis in GBM cells and in intracranial GBM models

RT-induced SAM elevation could either be due to increased SAM synthesis or diminished SAM consumption. To distinguish between these possibilities, we performed stable isotope tracing to directly measure SAM synthesis. This technique employs a heavily labeled methionine tracer containing 5 atoms of ^13^C instead of ^12^C (Fig. 2A). ^13^C_5_ methionine is non-radioactive but can be detected by mass spectrometry as it weighs 5 mass units more than ^12^C-containing methionine. Downstream metabolites newly formed from the methionine tracer such as SAM and SAH also weigh more than normal, which allows us to detect changes in metabolic activity post-RT (Fig. 2A). SAH gets converted to homocysteine which can then be recycled to methionine by gaining an unlabeled ^12^C from the folate cycle, generating recycled methionine. Recycled methionine with 4 atoms of ^13^C is then converted to recycled SAM by MAT2A (Fig. 2A).

**Figure 2.**
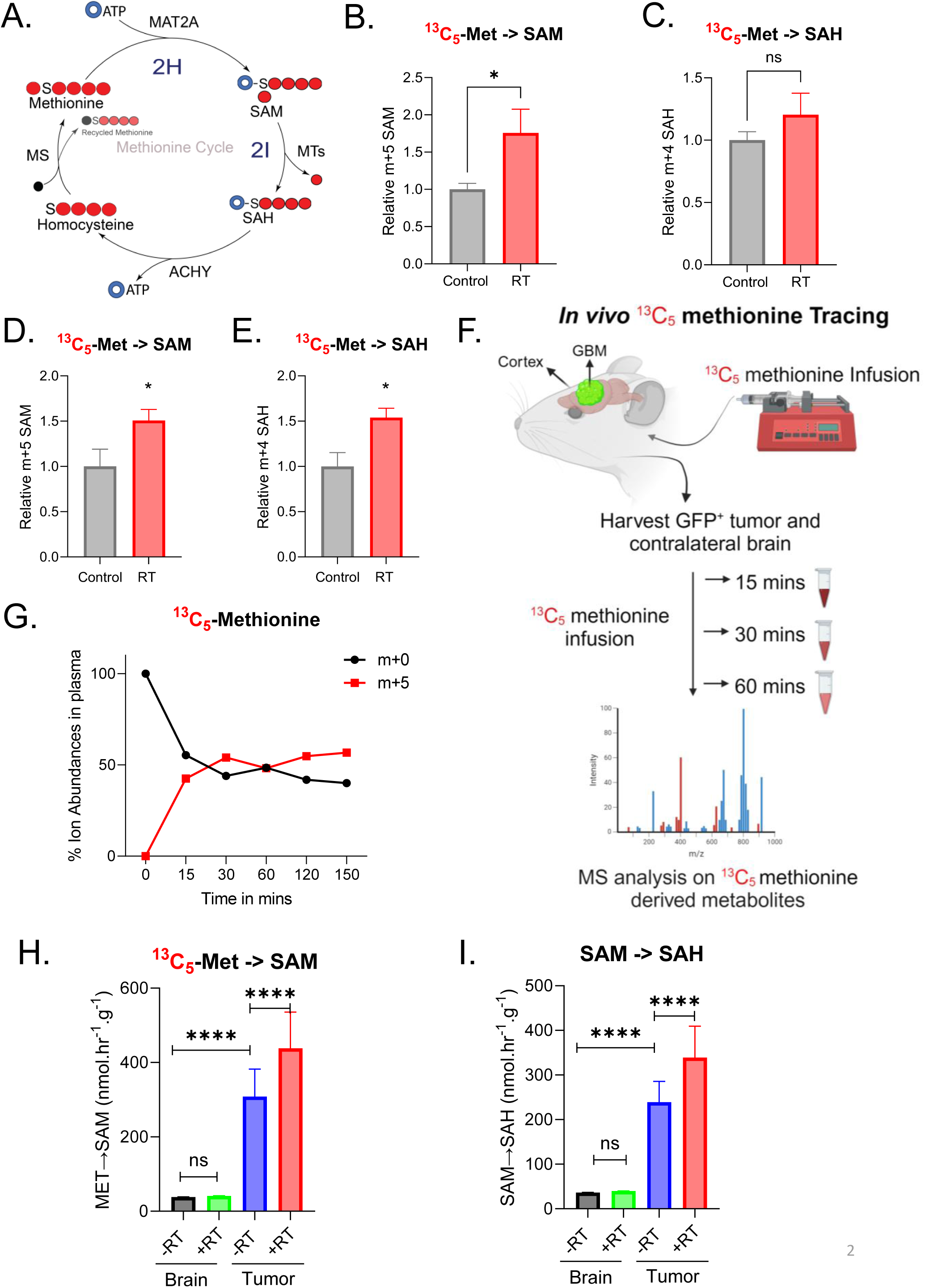
RT acutely increases SAM synthesis in GBM cells and in intracranial GBM models. A). Schema of ^13^C_5_ methionine carbons (red circles) redistribution into methionine cycle intermediates. The black circle indicates unlabeled ^12^C methionine. B-C). U87 cells were incubated with ^13^C_5_ methionine just before RT (8 Gy) and metabolites were harvested (1 hr post-RT) for LC/MS analysis. Relative m+5 SAM and m+4 SAH abundances are plotted. Data are represented as mean±SEM for 7 biological replicates. D-E). HF2303 spheres were incubated with ^13^C_5_ methionine just before irradiation (8 Gy) and metabolites were analyzed using LC-MS. Relative ion abundances of m+5 SAM and m+4 SAH are plotted. Data are represented as mean±SEM for 8 biological replicates. P values of SAM and SAH are 0.0423 and 0.0106 respectively. F). Schema of ^13^C_5_ methionine infusion into mouse models of brain cancer. ^13^C_5_ methionine was infused into HF2303 PDX- bearing mice. Mice were either untreated or treated (8 Gy) and serially euthanized at various time points (15, 30 and 60 mins) and the GBM tumor and cortex were harvested to measure metabolite abundances and estimate metabolic fluxes. G). Time course of m+5 methionine plasma enrichment in mice undergoing ^13^C_5_ methionine infusions. H-I). Absolute metabolic fluxes of SAM and SAH in normal brain and GBM with or without RT from N=3 biological replicates. A-I). *P ≤ 0.05 and ****P ≤ 0.0001.

In U87 cells, RT acutely increased the abundance of ^13^C_5_-labeled SAM (Fig. 2B) derived from ^13^C_5_ Methionine (Fig. S2A). Further, ^13^C_4_-labeled SAH did not decrease post-RT, indicating that elevated SAM levels is due to increased SAM synthesis from methionine by MAT2A, rather than decreased SAM consumption by methytransferase reactions (Fig. 2C). This is further evidenced by a significant increase in recycled SAM post-RT (Fig. S2B, m+4) and a trend towards increased recycled methionine (Fig S2C, m+4) post-RT. We confirmed our findings in a patient derived HF2303 neurosphere model, where we observed a significant increase in ^13^C_5_-labeled SAM and ^13^C_4_-labeled SAH levels post-RT (Fig. 2D,2E and Fig. S2D, S2E and S2F). In a third patient derived GBM6 explant model, we observed a trend towards increased SAM labeling after RT that did not reach statistical significance (Fig. S2G, S2H and S2I).

To understand the metabolic fates of methionine in brain tumors *in vivo*, we infused ^13^C_5_ methionine into the jugular vein of mice and followed methionine-derived ^13^C into its downstream fates in cortex and GBM tissue (Fig. 2F). To determine optimal methionine infusion conditions, we tested 3 different dose ranges (low, medium and high) of methionine (Fig. S2J). All three doses of methionine introduced ^13^C_5_ methionine into circulation, labeling downstream metabolites SAM and SAH in the mouse brain in a dose-dependent fashion indicating that circulating methionine is readily available for tissues (Fig. S2K and S2L). While the medium and high doses of methionine increased arterial methionine concentrations by 2-3-fold, low dose methionine infusions achieved nearly 50% labeling of circulating methionine without dramatically elevating arterial methionine concentrations (Fig. S2J-L). We therefore opted to use the low dose methionine to study cortical and tumor metabolism *in vivo*.

Using these infusion conditions, we irradiated HF2303-bearing mice with cranial RT (8 Gy) and immediately infused the mice with ^13^C_5_ methionine. The relative ^13^C_5_ methionine abundance in plasma reaches a steady state within 15 mins of infusion (Fig. 2G). We used fluorescent-guided microdissection to isolate GFP-expressing intracranial tumor from the surrounding cortex serially at 15mins, 30 mins and 60 mins post-RT (Fig. 2F). Following the entry into the tissues, ^13^C_5_ methionine is metabolized by the methionine cycle into downstream metabolites. Monitoring the ^13^C label incorporation into these metabolic pathways helped us determine the metabolic activity and metabolic flux through the methionine cycle. To estimate the rates of SAM synthesis, we calculated the metabolic fluxes of brain and tumor in both control and irradiated mice (Fig. S2N-O and Fig. 2H). The rate of SAM synthesis in the tumor is significantly higher than in the normal brain in unirradiated mice (Fig. 2H, blue vs. black). Interestingly, we found that while the SAM synthesis rate did not change with irradiation in normal brain, there is a further increase in SAM synthesis in brain tumors post-RT (Fig. 2H, red vs. blue). We also calculated the flux rates for label incorporation into SAH (Fig. 2I and S2P). We found that there is an increase in flux towards SAH production in the tumor compared to the normal brain. While RT did not change the SAH production rate in normal brain, there is a significant increase in SAH production in tumors post-RT (Fig. 2I). Interestingly, we found that tumors have significantly higher methionine recycling compared to cortex and is unchanged post-RT (Fig. S2M and S2Q). Thus, we conclude that RT increases the SAM synthesis rate in brain tumors, while the SAM synthesis rate remains unchanged following RT in the normal brain.

### Disrupting methionine metabolism enhances the responsiveness of GBMs to RT

We next sought to investigate whether elevated SAM synthesis in GBM drives RT resistance. We disrupted methionine-driven SAM synthesis by depriving cells of methionine or by pharmacologically inhibiting MAT2A, the enzyme responsible for synthesizing SAM from methionine (Fig. 2A). Methionine restriction sensitized both the immortalized U87 cell line and patient-derived HF2303 neurospheres to RT, with an enhancement ratio of 3.3 in U87 cells (Fig. 3A) and 1.8 in HF2303 cells (Fig. 3B). We observed similar results using the newly described blood-brain barrier penetrant inhibitor of MAT2A, AGI-41998.^25^ Treatment with AGI-41998 sensitized both U87 cells (Fig. 3C) and HF2303 (Fig. 3D) neurospheres to RT with enhancement ratios of 1.5 in both cases.

**Figure 3:**
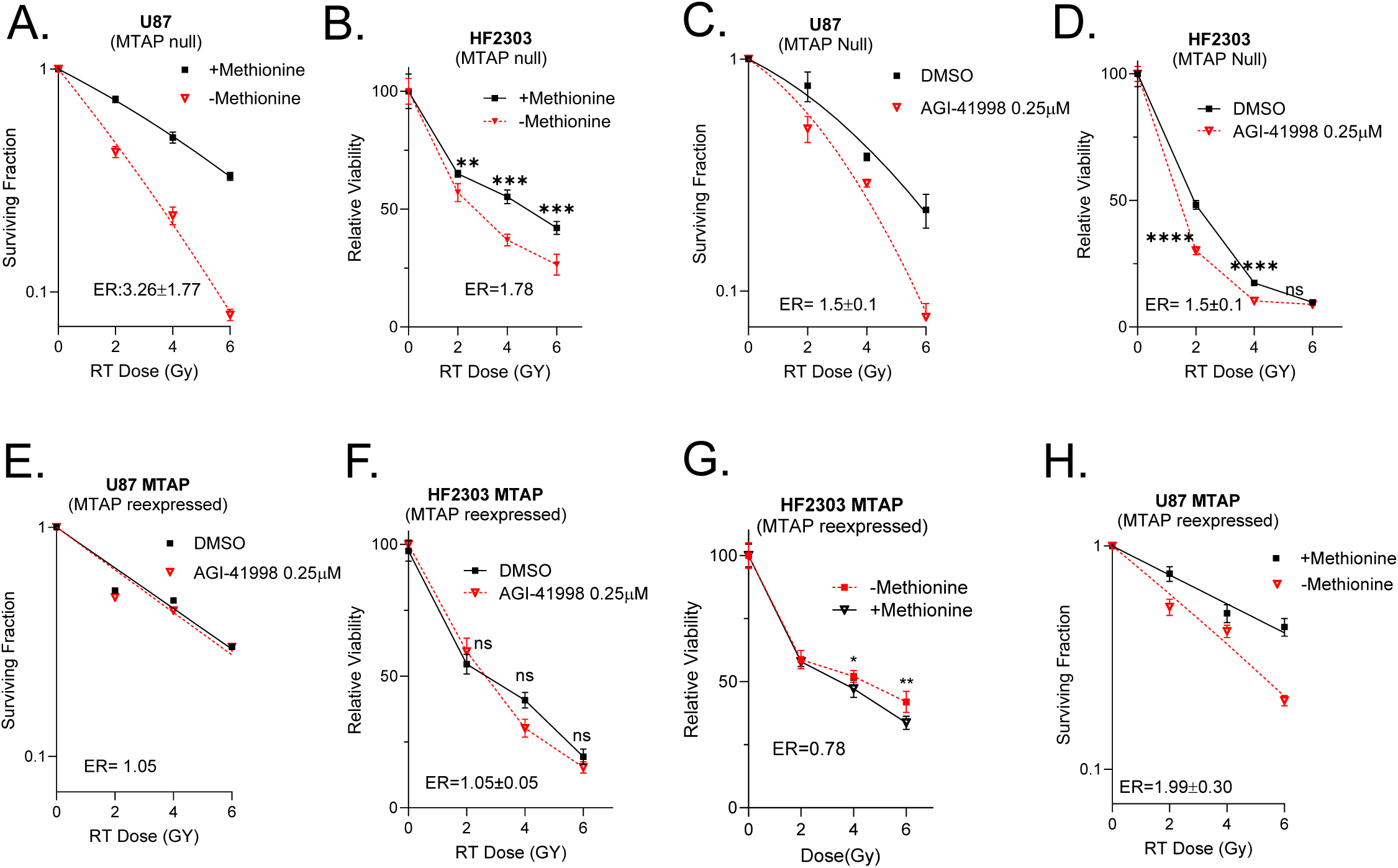
Disrupting methionine metabolism enhances the responsiveness of GBMs to RT. A). U87 cells were either were grown in media with methionine or media depleted of methionine and irradiated with indicated doses of RT followed by cell viability by clonogenic survival assay. Enhancement ratio (ER) is the ratio of DMID control and DMID methionine depleted cells. ER<1 indicates radioprotection and ER>1 indicates radiosensitization. Data is presented as mean±SEM for 3 biological replicates. B). HF2303 neurospheres were either were grown in media with methionine or media depleted of methionine and irradiated with indicated doses of RT followed by cell viability detected by cell titer glo kit 7-10 days after replating. ER is ratio of GI50 of control cells and GI50 of methionine depleted cells. C). U87 cells were either untreated or treated with AGI-41998 and irradiated with indicated doses followed by clonogenic survival assay. Enhancement ratio (ER) is the ratio of DMID control and DMID treated cells. Data is presented as mean±SEM for 3 biological replicates. D). HF2303 neurospheres were either untreated or treated with AGI-41998 and irradiated with indicated doses followed by long term sphere formation assay. ER is ratio of GI50 of control cells and GI50 of AGI-41998 treated cells. E-F). U87 MTAP and HF2303 MTAP spheres were either untreated or treated with AGI-41998 and irradiated with indicated doses followed by clonogenic survival assay and cell viability by cell titer glo kit 7-10 days after replating respectively. Data is presented as mean±SEM for 3 biological replicates. G-H). U87 MTAP or HF2303 MTAP neurospheres were either were grown in media with methionine or media depleted of methionine and irradiated with indicated doses of RT followed by clonogenic survival assay and long-term sphere formation assay respectively (3 biological replicates). A-H). *P ≤ 0.05, **P ≤ 0.01, ***P ≤ 0.001 and ****P ≤ 0.0001.

Both U87 cells and HF2303 cells harbor a deletion of *MTAP*, a methionine salvage enzyme that is deleted in ∼40% of GBM tumors. Cells with deleted *MTAP* have been found to be selectively sensitive to MAT2A inhibition, though there is some question whether this selectivity exists *in vivo*, due to an apparent lack of elevation of the metabolite MTA.^26^ However, when we interrogated xenograft GBM tissues and our own patient samples, we found approximately 3- fold increase in xenograft GBM tissue and 2-fold increase in MTA in GBM patients that lack MTAP expression (Fig. S3A-S3C). We thus re-expressed MTAP in both U87 cells (Fig. S3D) and HF2303 neurospheres (Fig. S3E) and re-assessed the relationship of methionine metabolism and RT response. Reexpression of MTAP completely reversed the ability of MAT2A inhibitors to sensitize GBM cells to RT (Fig. 3E and 3F). Similarly, MTAP re-expression reversed the ability of methionine restriction to sensitize HF2303 cells to RT and partially rescued this phenotype in U87 cells (Fig. 3G and 3H). Together, these results suggest that disrupting methionine metabolism might have greatest efficacy in GBMs lacking *MTAP*.

We also sought to determine if MAT2A inhibition also influenced the response to other genotoxic stressors. To this end, U87 cells treated with MAT2A inhibitor showed greater sensitivity to a genotoxic chemotherapy agent, Bleomycin^27^ (Fig. S3F and S3H). We also confirmed our findings in a *MTAP-*deleted pancreatic cell line KP4 (Fig. S3G and S3I). MAT2A inhibition increased the sensitivity to bleomycin, indicating the generalizability of MAT2A inhibition in improving sensitivity to genotoxic stress.

### Disrupting methionine metabolism slows DNA repair in GBM

Because RT primarily kills cells through unrepaired double-stranded DNA breaks, we reasoned that SAM depletion may sensitize GBMs to RT by slowing double-stranded break (DSB) repair. Consistent with this hypothesis, methionine depletion delayed double strand break repair, especially at later time points post-RT in U87 cells (Fig. 4A and Fig. S4A). We confirmed these findings in HF2303 neurospheres, where methionine depletion delayed DNA repair (Fig. 4B and Fig. S4B). We observed similar results using MAT2A inhibition. AGI-41998 treatment increased the γ-H2AX foci at various time points post-RT (6 and 24 hours) in U87 cells (Fig. 4C and Fig. S4C). We confirmed our findings in the HF2303 neurosphere model, where MAT2A inhibition also delayed DNA repair (Fig. 4D and Fig. S4D).

**Figure 4:**
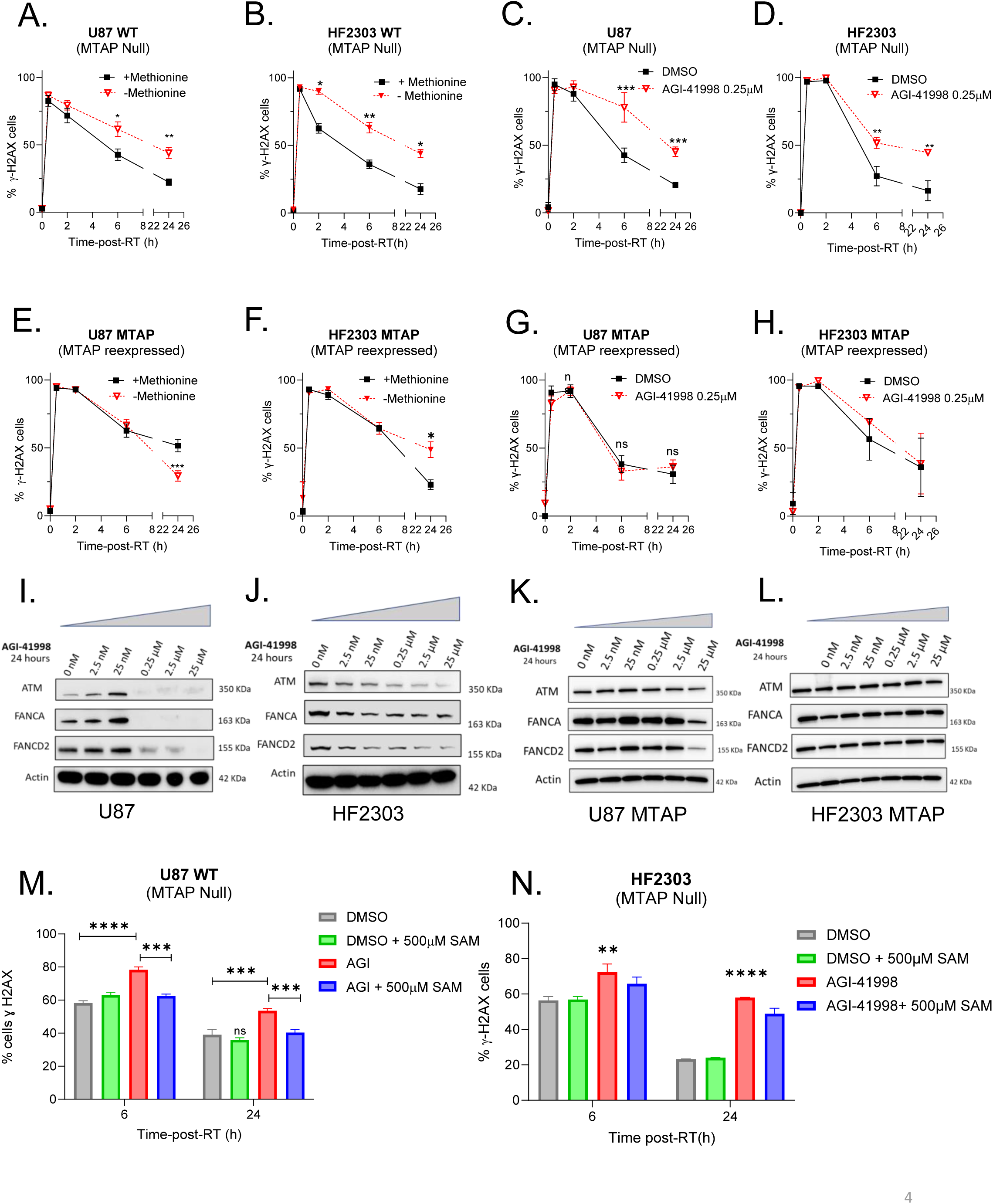
Disrupting methionine metabolism slows DNA repair in GBM. A-B). U87 cells or HF2303 spheres were grown in media with methionine or media depleted of methionine and irradiated (4 Gy). Cells or spheres were harvested at the indicated time points for γ-H2AX staining. Data is presented as mean±SEM for 3 biological replicates. C-D). U87 cells or HF2303 spheres were grown either untreated or treated with AGI- 41998, irradiated (4GY) and harvested for γ-H2AX staining at the indicated time points post-RT(4 Gy). Data is presented as mean±SEM for 3 biological replicates. E-F). U87 MTAP cells or HF2303 MTAP spheres were grown in media with methionine or media depleted of methionine and cells or spheres were harvested for γ-H2AX staining at the indicated time points post-RT(4 Gy). Data is presented as mean±SEM for 3 biological replicates. G-H). U87 MTAP cells or HF2303 MTAP spheres were untreated or treated with AGI-41998 and harvested for γ-H2AX staining at the indicated time points post- RT(4 Gy). Data is presented as mean±SEM for 3 biological replicates. I-J). Western blot showing the reduction of DNA repair proteins ATM, FANCA and FANCD2 in U87 cells and HF2303 neurospheres respectively. K-L). Western blot showing the levels of DNA repair proteins ATM, FANCA and FANCD2 in U87 MTAP cells and HF2303 MTAP neurospheres respectively. M-N). U87 cells and HF2303 neurospheres were treated with SAM overnight and retreated with SAM 1 hr before RT (4 Gy) and collected at indicated time points for γ-H2AX staining. Data is presented as mean±SEM for 3 biological replicates. N). P values were obtained in comparison with the control (DMSO). A-N). *P ≤ 0.05, **P ≤ 0.01, ***P ≤ 0.001 and ****P ≤ 0.0001.

We next assessed whether the expression of *MTAP* influenced the ability of SAM inhibition to regulate DSB repair. Forced re-expression of MTAP in U87 cells (Fig. 4E and Fig. S4E) and HF2303 neurospheres (Fig. 4F and Fig. S4F) partially rescues the delay in DNA repair caused by methionine restriction. Similarly, re-expression of MTAP reversed the ability of the MAT2A inhibitor AGI-41998 to slow DNA repair in both U87 cells (Fig. 4G and Fig. S4G) and HF2303 neurospheres (Fig. 4H and Fig. S4H).

We then sought to understand the mechanism by which inhibiting SAM synthesis regulated the RT response and DNA repair. In other contexts, MAT2A inhibition alters splicing efficiency and affects the levels of numerous proteins, including some involved in the DNA damage response.^28^ We found that MAT2A inhibition reduces the DNA repair proteins such as ATM, FANCA and FANCD2 in MTAP-deleted U87 cells and HF2303 neurospheres in a concentration dependent fashion (Fig. 4I and 4J). Re-expression of MTAP in U87 cells and HF2303 neurospheres rescued the AGI-41998 induced reduction of DNA repair proteins (Fig. 4K and 4L). Similarly, methionine depletion reduced the DNA repair proteins in MTAP null cells, while re-expression of MTAP rescued the reduction of DNA repair proteins (Fig. S4I). Based on these results, we conclude that radiosensitization induced by inhibiting SAM production in MTAP-deleted cells is due to delayed DNA repair due to decreased expression of proteins involved in the DNA damage response.

To further support this conclusion, we asked if SAM supplementation could rescue the AGI-41998-mediated reduction in DNA repair proteins and delay in DNA repair. To this end, we supplemented U87 cells with SAM, which rescued the reduction of DNA repair proteins (Fig. S4J). Similarly, we supplemented U87 cells with SAM and evaluated DNA repair. Consistent with our previous results, MAT2A inhibition delayed DNA repair at 6 and 24 hours. SAM supplementation rescued the delay in DNA repair by the MAT2A inhibitor, indicating the delayed DNA repair is due to the depletion of SAM levels by the MAT2A inhibitor (Fig. 4M and Fig. S4K). We confirmed these results in a neurosphere HF2303 model, where SAM supplementation reversed the AGI-41998- mediated delay DNA repair (Fig. 4N and Fig. S4L).

### MAT2A inhibition radiosensitizes *MTAP*-deleted flank models of GBM

We then sought to understand if inhibiting SAM synthesis can overcome GBM RT resistance in tumor models *in vivo*. We first evaluated the ability of the MAT2A inhibitor AGI-41998 to reduce SAM levels *in vivo*. We treated mice carrying flank tumors with two different doses of the drug (Fig. S5A). We found that administering AGI-41998 at 30 mg/kg and 60 mg/kg led to detectable and dose-dependent drug concentrations in the plasma, flank tumor, and normal brain. Furthermore, both doses of drug significantly reduced SAM levels in plasma, flank tumor, and brain (Fig. S5B). Based on these findings, we chose to use 30 mg/kg of AGI-41998 for *in vivo* experiments, since both 30 mg/kg and 60 mg/kg achieved similar levels of SAM depletion.

We then utilized patient-derived, MTAP-deleted, and RT-resistant models of GBM (GBM6 and GBM38), from the Mayo Clinic repository to generate flank tumors in mice (Fig. 5A). Once the flank tumors reached 80-100mm^3^, the mice were randomized into 4 groups: Control, RT alone, AGI-41998, or AGI-41998 +RT. We analyzed a subset of these tumors 2 hours post-RT using LC-MS (Fig. 5A). In both GBM38 and GBM6 models, AGI-41998 (30 mg/kg) significantly reduced the SAM levels when used as monotherapy or in combination with RT (Fig. 5B and 5C). In both GBM6 and GBM38 flank tumor models, AGI-41998 treatment alone modestly slowed tumor growth (Fig. 5D and 5E). RT also modestly reduced tumor growth while the combination significantly slowed tumor growth and significantly increased the time to tumor tripling (Fig. 5F and 5G). Median days to tumor tripling are 6(Control), 9(AGI-41998), 9(RT) and 24(AGI- 41998+RT) for GBM6, 11(Control), 17(AGI-41998), 20(RT), 37.5(AGI-41998+RT) for GBM38 respectively. An unchanged body weight in both the tumor models indicates minimal effects of treatment on normal tissues (Fig. S5C and S5D). We also assessed SAM levels in the endpoint GBM6 and GBM38 flank tumors (Fig. S5E and S5F). AGI- 41998 treatment alone or in combination with RT significantly reduced SAM levels in end point tumors, indicating the improved efficacy of the combination treatment is the result of MAT2A inhibition along with RT.

**Figure 5:**
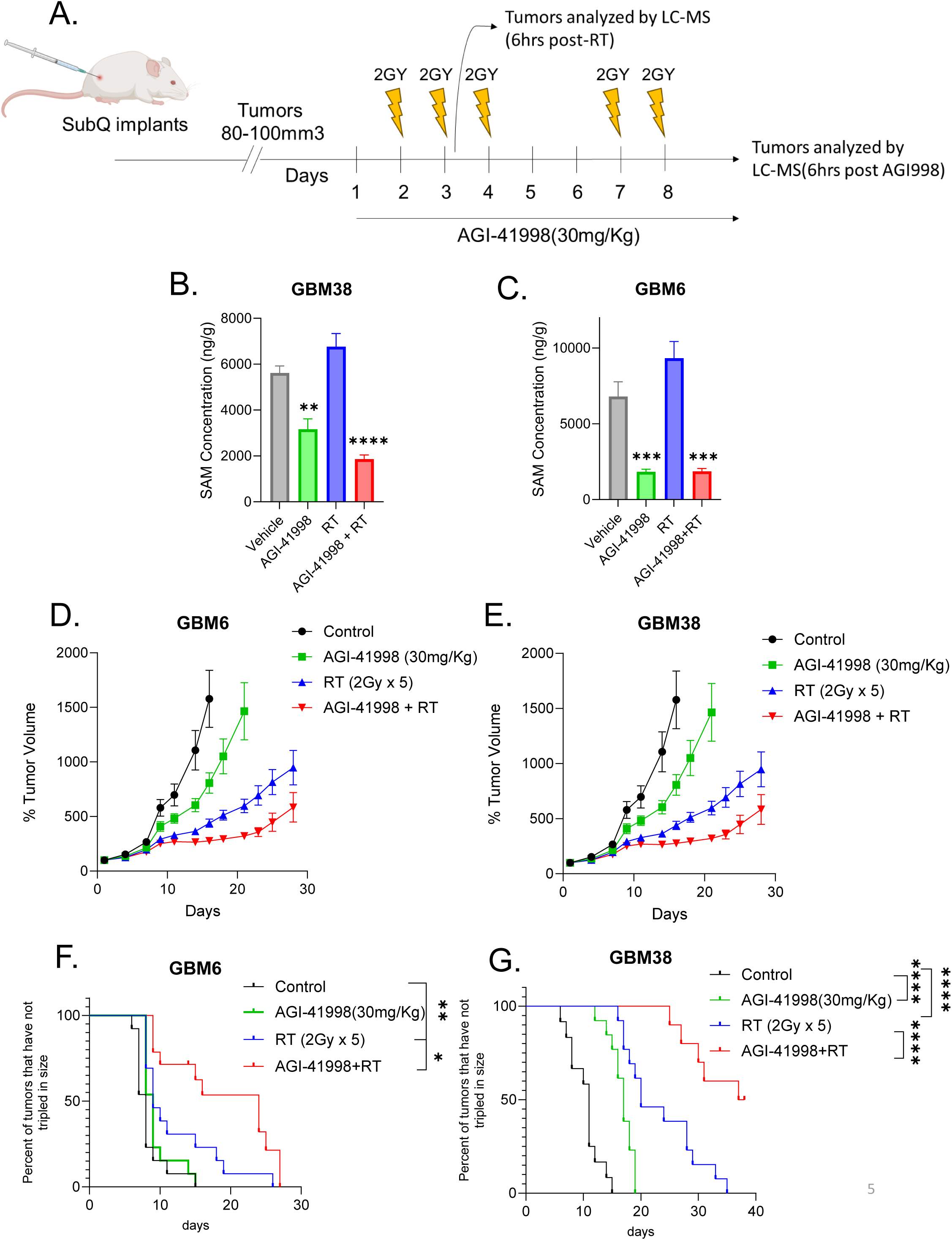
MAT2A inhibition radiosensitizes *MTAP*-deleted flank models of GBM. A). Schematic timeline of GBM6 and GBM38 flak tumor models. GBM6 and GBM38 xenograft GBM models were established and randomized into 4 groups, Control, AGI- 41998, RT, or AGI-41998+RT. AGI-41998(30 mg/Kg) was administered via oral gavage daily starting one day prior to RT and continued till end point. AGI-41998 was given 1 hour before RT(2 Gy) over six days on weekdays. B-C). GBM6 or GBM38 PDXs were implanted into flanks of mice (Fig. 5A). Mice were randomized and were given AGI- 41998(30 mg/Kg) daily via oral gavage and 2 fractions of RT (2 Gy) and tumors were harvested 6 hours post last dose of RT and were analyzed by LC-MS analysis. P values were obtained in comparison with the control (Vehicle). D-E). Tumor volumes of each treatment group is normalized to tumor volume of day1 of treatment. Data is presented as mean±SEM from 13-14 tumors from 7 mice per group. F-G). Kaplan-Meier curves of time to tumor tripling. P values for Control Vs. AGI-41998 are 0.1405 Fig. F and <0.0001 Fig. G, Control Vs. RT are 0.0088 Fig. F and <0.0001 Fig. G, RT Vs AGI-41998+RT are 0.0170 Fig. F and 0.0006 Fig. G. F- P values were calculated using Log-rank (Mantel- Cox) test. A-G). *P ≤ 0.05, **P ≤ 0.01, ***P ≤ 0.001 and ****P ≤ 0.0001.

### MAT2A inhibition radiosensitizes intracranial models of GBM

Since the GBM microenvironment and the blood-brain barrier can greatly influence tumor biology and drug efficacy, we next sought to determine if AGI-41998 can overcome GBM RT resistance in *intracranial* models of GBM. We utilized patient-derived RT-resistant luciferase expressing HF2303 neurospheres to perform intracranial injections in mice. Once the tumors were detectable by bioluminescence, we randomized the mice into 4 groups, Control, RT, AGI-41998, or AGI-41998+RT (Fig. 6A). We analyzed a subset of these tumors 1 hour post-RT using LC-MS. We found that 30 mg/Kg of AGI-41998 significantly reduced intracranial SAM levels (Fig. 6B) and increased levels of the MAT2A precursor methionine (Fig. S6A), thereby severely depleting the SAM/Methionine ratio (Fig. 6C), which indicates strong target engagement in intracranial GBM. SAH levels were not dramatically different in AGI-41998 -treated tumors (Fig. S6B).

**Figure 6:**
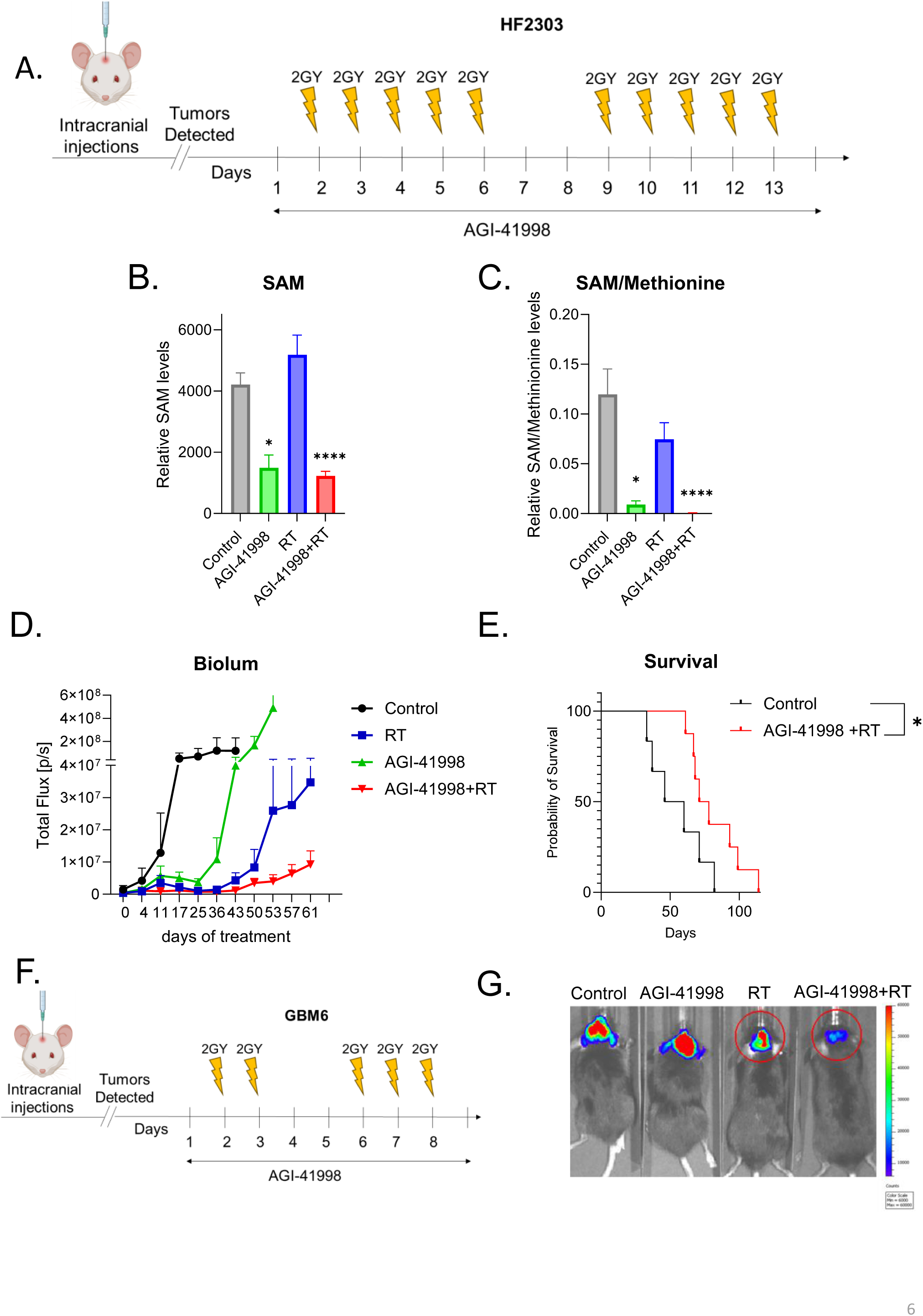
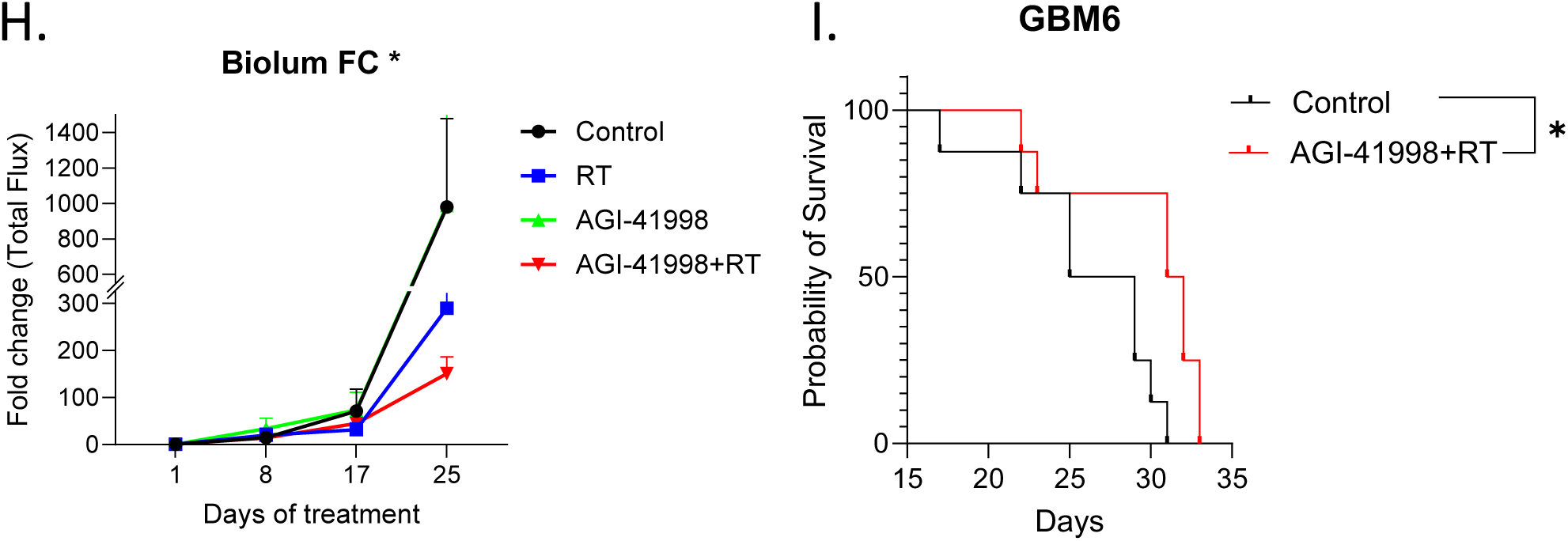
MAT2A inhibition radiosensitizes intracranial models of GBM. A). Schematic timeline of HF2303 intracranial tumor model. Luciferase positive HF2303 PDXs were orthotopically injected and once they were detectable by bioluminescent imaging the tumor-bearing mice were randomized into 4 groups, Control, AGI-41998, RT, or AGI-41998+RT (6-8 mice per group). AGI-41998 (30 mg/Kg) was given via oral gavage daily starting one day prior RT and ending one day after RT. AGI-41998 was given 1 hour before RT (2 Gy) over ten days on weekdays. B-C). HF2303 tumor bearing mice were randomized into 4 groups. Mice were given AGI-41998 (30 mg/Kg) oral gavage and one dose of RT (8 Gy) and harvested 1 hour after RT. Data is represented as mean ± SEM from 15 (Control, RT and AGI-41998+RT) to four (AGI-41998) data points. P values were obtained in comparison to the control. D). Total flux of equal Region Of Interests (ROIs) at each time point are presented as mean ± SEM from six (Control) to eight (AGI-41998, RT and AGI-41998+RT) mice per group. E). Kaplan-Meier survival curves. P values of Control Vs. AGI-41998+RT is 0.04, P values were calculated using Log-rank (Mantel-Cox) test. F). Schematic timeline of GBM6 intracranial tumor model. Luciferase positive GBM6 PDXs were orthotopically injected and the mice were randomized into 4 groups, Control, AGI-41998, RT, or AGI-41998+RT (6-8 mice per group). G). Representative pictures of tumors after luciferin was injected to the mice. H). Total flux for each treatment group with same ROIs is normalized to the total flux on day 1 to monitor the tumor growth. Data is represented as mean ± SEM from six (AGI-41998) to eight (Control, RT and AGI-41998+RT) I). Kaplan-Meier survival curves of GBM6 tumor bearing mice. P values for Control Vs. AGI-41998+RT are 0.01. P values were calculated using Log-rank (Mantel-Cox) test. A-I). *P ≤ 0.05 and ****P ≤ 0.0001.

While AGI-41998 monotherapy and RT alone only modestly reduced tumor growth, AGI-41998 in combination with RT profoundly reduced intracranial tumor growth (Fig. 6D). Consistent with these findings, mice treated with AGI-41998 in combination with RT had a median survival of 74.5 days, which was significantly improved compared to control mice, which had a median survival of 53 days (Fig. 6E). Neither RT alone nor AGI-41998 alone significantly improved survival compared to control (Fig. S6C).

To extend our findings to other GBM models, we used the luciferase-expressing MTAP-deleted GBM6 RT-resistant model to perform intracranial injections in mice. Once the tumors were detectable by bioluminescent imaging, we randomized the mice into 4 groups, Control, RT, AGI-41998 (15mg/kg), or AGI-41998+RT (Fig. 6F). AGI-41998 alone did not reduce tumor growth, RT alone slightly reduced tumor growth, and this effect was greater in the combination group (Fig. 6G and 6H). Consistent with these observations, AGI-41998 in combination with RT (median survival 31.5 days) significantly improved the survival of mice compared to controls (median survival 27 days, Fig. 6I). Monotherapy with either AGI-41998 or RT had minimal survival benefits (Fig. S6D). AGI-41998 by itself or in combination with RT caused slight weight loss in mice in both intracranial tumor studies (Fig. S6E and S6F). Thus, inhibiting MAT2A can potentiate the efficacy of RT in GBM *in vivo*, but does not appear to cure tumors.

We further investigated if depleting methionine levels (as opposed to inhibiting MAT2A) can improve RT efficacy, as it did *in vitro*. To this end, we utilized methioninase, a engineered human methionine degrading enzyme, to deplete serum methionine levels.^29^ We intracranially injected GFP and luciferase expressing GBM6 cells into mice and once the tumors were detectable by bioluminescent imaging, we randomized the mice into 4 groups, Control, RT (2 Gy*5), methioninase (100mg/Kg), and methioninase+RT (Fig. S6G). In contrast to MAT2A inhibition, methioninase treatment combined with RT did not reduce tumor growth and did not have any survival benefit (Fig. S6H and S6I). Methioninase treatment was well tolerated in mice with minimal weight loss (Fig. S6J). We further analyzed the end point tumors using LC-MS analysis. Methioninase treatment significantly reduced the methionine levels in plasma, brain, and tumors (Fig. S6K). However, the depletion of methionine failed to reduce the SAM levels in plasma, brain, and tumor (Fig. S6L), possibly explaining the lack of efficacy and survival benefit of methioninase treatment along with RT.

We also asked if dietary methionine restriction improved RT efficacy in GBM. Unfortunately, methionine diet restriction along with RT did not reduce tumor growth and did not have any survival benefit in orthotopic GBM38 tumors (Fig. S6M). Analysis of the endpoint tumors further revealed that while methionine diet restriction reduced methionine levels in intracranial tumors, it failed to reduce SAM levels (Fig. S6N), potentially explaining the lack of efficacy of methionine diet restriction with RT. Based on these results, we conclude that depletion of SAM in intracranial GBMs can augment the effects of RT, and that this is more feasibly achieved by MAT2A inhibition than with lowering of circulating methionine levels.

### RT-dependent SAM synthesis is an active signaling event

Lastly, we sought to understand how RT activates SAM synthesis (Fig. 1-2). We reasoned that this RT-induced metabolic activation could either be due to an active signaling event, or a reflexive activation due simply to increased SAM demand following DNA damage. The rate-limiting enzyme in SAM synthesis is MAT2A, which is regulated through numerous mechanisms including post-translational phosphorylation.^30^ Using phosphorylation-specific gel electrophoresis, we identified an increase in a slower migrating MAT2A band following RT, consistent with RT-induced phosphorylation of MAT2A (Fig. 7A, leftmost two columns in phosphogel). To identify the signaling pathways responsible for this RT-induced phosphorylation, we tested a panel of pharmacologic inhibitors of several kinases involved in the DNA damage response including ATM, ATR, DNA-PK, mTOR and MEK (reported to interact with MAT2A).^30,31^ Inhibition of ATR and MEK, (but not ATM, DNA-PK, or mTOR) reduced the RT-induced phosphorylation of MAT2A (Fig. 7A). We confirmed these findings in two other models, HF2303 and GBM6, where RT-induced phosphorylation of MAT2A and is blocked by ATR or MEK inhibition (Fig. 7B and 7C).

**Figure 7:**
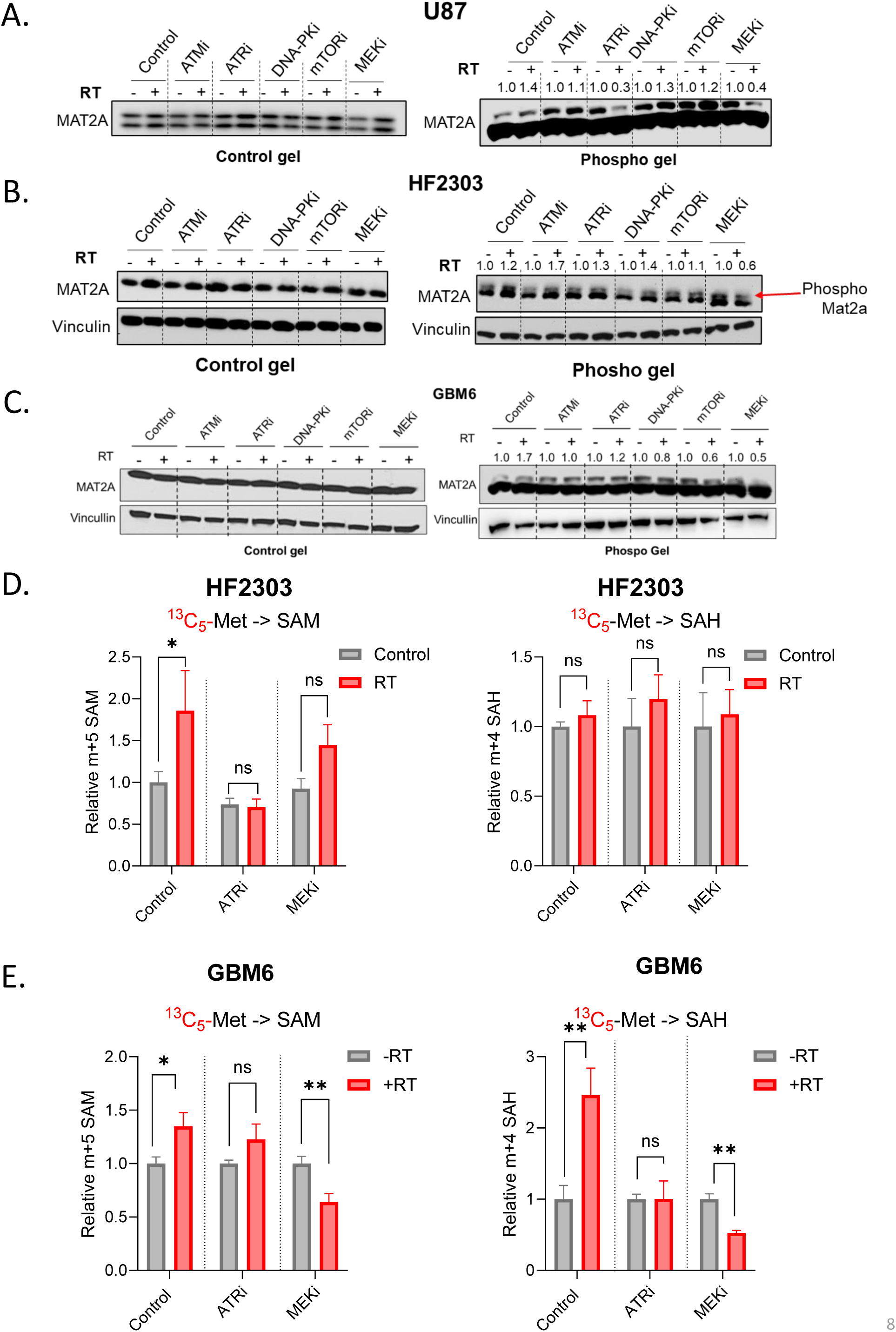
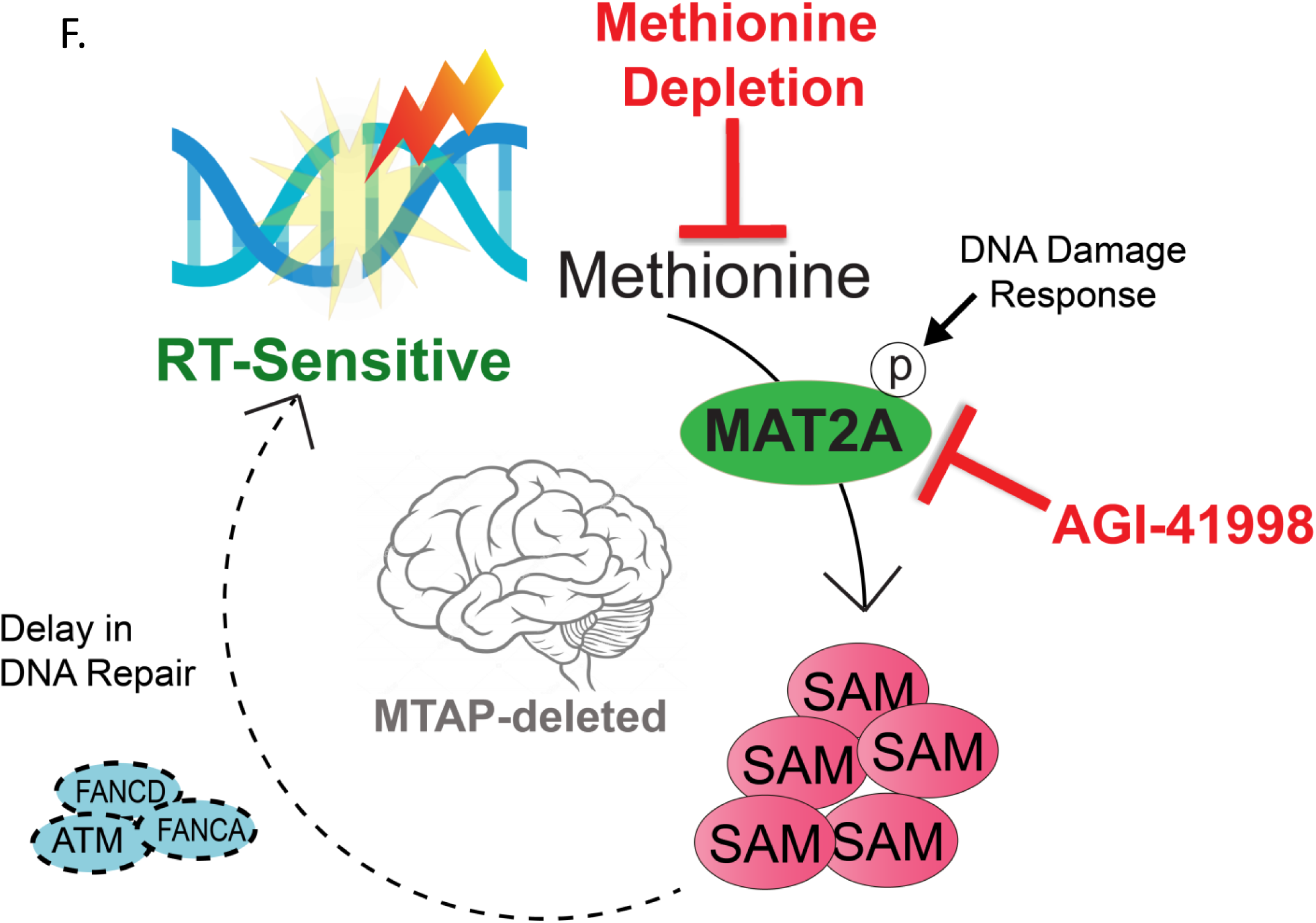
RT-dependent SAM synthesis is an active signaling event. U87 cells (A) or HF2303 and GBM6 spheres (B-C) were either untreated or treated with indicated inhibitors for 1 hour and sham irradiated or irradiated (8 Gy) and harvested 1 hour post-RT. Proteins were separated by gel electrophoresis on SuperSep control or Phos-tag gels and blotted using indicated antibodies. (D-E). HF2303 or GBM6 spheres were either untreated or treated with indicated inhibitors for 1hour. Then ^13^C_5_ methionine was added to cultures and they were immediately either untreated or irradiated (8 Gy). One hour later, spheres were harvested and the conversion of methionine into SAM and SAH was determined by LC-MS analysis. *p < 0.05 and **p < 0.01, compared with the control. F). A schematic summary

Based on these results, we asked if RT-induced phosphorylation of MAT2A is causal for SAM synthesis and if we can block it using inhibitors of ATR and MEK. We pretreated the cells for 1 hour before RT and performed stable isotope tracing to assess the SAM synthesis (Fig. 7D). In HF2303 spheres, RT acutely induces SAM synthesis and is consistent with our previous data. RT-induced synthesis of SAM is blocked by ATR and MEK inhibitors. We confirmed our findings in another patient-derived GBM model, GBM6 where RT-induced synthesis of SAM and SAH is blocked by both ATR and MEK inhibitors (Fig. 7E). Our results suggest that RT-induced phosphorylation of MAT2A boosts its activity, increasing SAM synthesis and can be blocked by ATR or MEK inhibition.

## Discussion

In this work, we have made the first direct measurements of methionine metabolism in brain tumors *in vivo* and defined links between altered methionine metabolism, DNA repair and treatment resistance in GBM. Compared to the cortex, GBM tumors have elevated rates of methionine-derived SAM synthesis, which increase further acutely following RT. RT induces the phosphorylation of MAT2A, the rate limiting enzyme in SAM synthesis and inhibiting RT-induced ATR and MEK signaling blocks both RT-induced MAT2A phosphorylation and the activation of SAM synthesis by RT. Thus, the ability of RT to activate methionine-driven SAM synthesis appears to be an active signaling event mediated by the DNA damage response. Consistent with this hypothesis, blocking RT-induced SAM synthesis *in vitro* through methionine depletion or MAT2A inhibition slows DNA repair and helps overcome GBM RT resistance. Mechanistically, SAM depletion reduces the expression of DNA repair proteins ATM, FANCA and FANCD2, likely due to the high levels of SAM needed to facilitate efficient mRNA splicing and subsequent translation of these proteins.^28^ These effects are most pronounced in GBM models that lack the methionine-salvage enzyme *MTAP* and are reversed by *MTAP* re-expression, which suggests that this therapeutic strategy will be most effective in *MTAP-*deleted GBM and may have minimal normal tissue toxicity. *In vivo*, MAT2A inhibition depletes SAM levels, delays tumor growth and radiosensitizes flank and intracranial GBM models. Depleting circulating methionine *in vivo* had minimal therapeutic effects, likely because this strategy failed to deplete SAM in tumors. These findings implicate RT-induced increase in SAM synthesis as a key link between methionine metabolism and DNA repair that can be targeted using MAT2A inhibitors.

These findings contribute to the existing body of research showing that disrupting methionine metabolism can enhance the susceptibility of cancers to genotoxic stress.^18,32–35^ For instance, limiting methionine improved the response of colorectal cancer and RAS-driven sarcoma mouse models to 5-flurouracil treatment to RT respectively, in part due to alterations in nucleotide and antioxidant metabolism^18^. Similarly, a small clinical trial suggested that methionine depletion may synergize with 5- fluorouracil in inhibiting thymidylate synthase activity in gastric cancers, again implicating folate metabolism.^36^ Our findings add to these studies by showing that targeting methionine metabolism synergizes with RT optimally when it depletes SAM and when GBMs lack MTAP. These findings implicate SAM-driven transmethylation as likely pathways that mediate GBM DNA repair and RT resistance. Depletion of SAM through MAT2A inhibition can disrupt RNA splicing, leading to reduced levels of DNA repair proteins such as the FANC family and ATM.^28^ Consistent with these findings, we observe that SAM depletion selectively reduces DNA repair proteins including FANCA, FANCD2 and ATM in *MTAP*-deleted GBMs and was sufficient to overcome GBM RT-resistance. There may be other methylation events driven by SAM/MAT2A that are crucial for DNA damage repair, but further studies are needed.

While methionine depletion or SAM inhibition both radiosensitized *in vitro* models of GBM, simply lowering methionine levels failed to reduce SAM levels and radiosensitize intracranial models of GBM. This might be attributed to the efficiency of the MAT2A enzyme, which may have a tight enough methionine binding efficiency to allow continued full SAM synthesis even when methionine levels drop by 50-70%.^37^ Our work suggests that investigators should ensure that methionine directed therapies deplete SAM if attempting to use them to augment RT efficacy in GBM. In our *in vivo* studies using methioninase and the methionine-restricted diet, tumor methionine levels dropped by 50-75%. It is possible that more severe methionine restriction could lower SAM levels sufficiently to synergize with RT.

Our studies suggest that MAT2A inhibitors, which are currently in clinical trials, may be useful for patients with GBM.^38^ Although MAT2A inhibitors were developed to target *MTAP*-deleted cancers selectively, there is some debate about whether this approach would be effective in GBM^26^. MTAP-deleted cancers appear to be sensitive to MAT2A inhibition due to the accumulation of the metabolite MTA, which increases the dependency on MAT2A-generated SAM. However, there is an apparent discrepancy between *in vitro* models of *MTAP*-deleted GBM, where the metabolite MTA accumulates to high levels and *in vivo* GBM tumors where MTA accumulation appears to be lower.^26^ Our data indicates that MTA levels are 3-5-fold higher in *MTAP*-deleted GBMs compared to contralateral brain tissue. Because re-expression of *MTAP* prevented the ability of MAT2A inhibition and methionine restriction to sensitize glioma models to RT, we speculate that these methionine-directed therapies will be most effective in patients whose gliomas lack *MTAP* and have the highest accumulation of MTA. Evaluating both *MTAP* expression and the amount of MTA elevation might be useful clinical correlates should this treatment strategy move to clinical trials.

Our work also suggests a mechanism that adaptively regulates SAM synthesis in response to DNA damage. Several regulatory mechanisms for MAT2A-driven SAM synthesis have been described. Phosphorylation of MAT2A stabilizes MAT2A and promotes SAM synthesis in liver cells.^30^ Acetylation of MAT2A has been shown to degrade MAT2A in hepatocellular cancer.^39^ Long ncRNA can transcriptionally regulate MAT2A expression post- RT.^40^ Tumor-initiating cells can alter methionine cycle activity, promoting tumor growth and resistance to therapy.^41,42^ Our data indicates acute changes in tumor MAT2A activity post RT via post translational modification leading to an increase in SAM synthesis adding to the numerous ways that the cell activates metabolic pathways after RT to help fuel survival.^43,44^

This study brings up several unanswered questions that require further exploration. While we know that RT induces SAM synthesis by phosphorylating MAT2A in a MEK-and ATR-dependent fashion, the exact mechanisms behind this regulation and precise phosphorylation sites are not yet certain. Additionally, there are likely additional RT-induced methylation events that regulate DNA repair and treatment resistance beyond the regulation of splicing and histone methylation. These remain to be explored. Lastly, and most importantly, there are numerous questions about how to translate these findings to the clinic. Our preclinical studies indicate that there is a significant increase in MTA accumulation in tumors compared to the contralateral cortex in MTAP-deleted models of GBM. It is currently unknown whether this metabolic alteration specific to GBM is dependent on MTAP deletion. Previous studies have suggested that the increase in MTA in human tumors is minimal, which may indicate that inhibiting MAT2A could be ineffective, even for *MTAP*-deleted tumors.^26^ Lastly, we utilized a recently developed blood-brain barrier permeable MAT2A inhibitor for this work, which achieved excellent SAM depletion intracranially.^25^ Whether the MAT2A inhibitors currently in clinical use effectively deplete SAM in brain tumors is not known and should be formally evaluated.

In summary, we have discovered a new regulatory link between DNA damage, SAM synthesis, and DNA repair. Disrupting this regulation can slow DNA repair and potentiate the effects of RT in *MTAP*-deleted brain cancers and may hold promise as a therapeutic intervention in this deadly disease.

## MATERIALS AND METHODS

### Cell culture and reagents

The U87 cell line was obtained from ATCC. HF2303 spheres were a kind gift from Dr. Alnawaz Rehemtulla. GBM6 and GBM38 xenografts were obtained from Mayo Clinic PDX national resource. The cell lines were authenticated by their respective repositories and were immediately used upon receipt. If cell lines were used for more than a year, they were reauthenticated using short tandem repeat profiling at the University of Michigan Advanced Genomics Core. Mycoplasma was tested (Cat# LT07-418, Lonza) monthly in our lab. U87 cells were cultured in DMEM (Cat# 11965-092, Gibco) with 10% FBS (Cat# S11550, Atlanta biologicals) and 1% PS (Cat# 10378-016, Gibco). HF2303 sphere were grown in DMEM F12 (Cat# 10565-018, Gibco) with B-27 supplement (Cat# 17504-044, Thermofisher), N2 supplement (Cat# 17502-048, Thermofisher), 100 μg/mL Normocin (Cat# ant-nr-1, InvivoGen), 100 U/mL PS (Cat# 15140122, Thermofisher), 20 ng/mL Epidermal growth factor (Cat# AF-100-15, PreproTech) and 20ng/mL Fibroblast growth factor (Cat# 100-18B, PeproTech). L-Methionine (^13^C_5_, 99%) was obtained from Cambridge Isotope Laboratories (Cat# CLM-893-H-PK). AGI-41998 for both *in vitro* and *in vivo* experiments was obtained from Agios pharmaceuticals. S-adenosylmethionine (Cat# B9003S, New England BioLabs), L-Methionine (Cat# J61904-18, Thermofisher).

Inhibitors: KU-60019, (S1570, Selleckchem), M3814 (S8586, Selleckchem), AZD6738 (S7693, Selleckchem), Rapamycin (AY-22989, Selleckchem), Mirdametinib (PD0325901, Selleckchem)

### Stable isotope tracing

U87 cells were labeled by culturing in DMEM media lacking glutamine, methionine, and cystine (Cat# 21013024, Thermo Fisher). The media was supplemented with 10% FBS and 1% PS, heavily labeled methionine (^13^C_5_ methionine, Cat# CLM-893-H-PK, Cambridge Isotope Laboratories), and the unlabeled glutamine and cystine. HF2303 spheres were cultures in DMEM/Nutrient Mixture F-12 Ham (Cat# D9785, Sigma-Aldrich) lacking L-glutamine, L-leucine, L-lysine, L-methionine, CaCl2, MgCl2, MgSO4, sodium bicarbonate, and phenol red. The media was supplemented with B-27 supplement (Cat# 17504-044, Thermofisher), N2 supplement (Cat# 17502- 048, Thermofisher), 100 μg/mL Normocin (Cat# ant-nr-1, InvivoGen), 100 U/mL PS (Cat# 15140122, Thermofisher), 20 ng/mL EGF (Cat# AF-100-15, PreproTech) and 20ng/mL EGF (Cat# 100-18B, PeproTech), heavy labeled methionine (^13^C_5_ methionine, Cat# CLM-893-H-PK, Cambridge Isotope Laboratories) and reconstituted with the remaining unlabeled amino acids and reagents. U87 cells or HF2303 spheres were plated a density of 3 million cells per 10 cm dish. Cells or spheres were treated with DMSO or AGI-41998 overnight and the media was changed to heavy labeled media, just before RT. The cells were harvested 1 hour after RT(8 Gy) to be processed by LC-MS analysis.

### Mass Spectrometry analysis

Cells or spheres were harvested and washed with 150mM Ammonium acetate and quenched using 80% ice-cold methanol. Flash- frozen tissues were homogenized in 80% ice-cold methanol. 100% ice-cold methanol was added to plasma samples to get to a final concentration of 80% methanol. The insoluble material was separated by centrifugation at >3000 rpm for 10mins at 4°c. The soluble metabolites were dried using Nitrogen purging. The dried metabolites were reconstituted and analyzed using mass spectrometry.

Ion Pairing Reverse phase LC-MS analysis: Analysis was performed on an Agilent system consisting of an Infinity Lab II UPLC coupled with a 6545 QTof mass spectrometer (Agilent Technologies, Santa Clara, CA) using a JetStream ESI source in negative mode. The following source parameters were used: Gas Temp: 250°C, Gas Flow: 13 L/min, Nebulizer: 35 psi, Sheath Gas Temp: 325°C, Sheath Gas Flow: 12 L/min, Capillary: 3500 V, Nozzle Voltage: 1500 V.

The UPLC was equipped with a 10-port valve configured to allow the column to be either eluted to the mass spectrometer or back-flushed to waste. The chromatographic separation was performed on an Agilent ZORBAX RRHD Extend 80Å C18, 2.1 × 150 mm, 1.8 μm column with an Agilent ZORBAX SB-C8, 2.1 mm × 30 mm, 3.5 μm guard column. The column temperature was 35°C. Mobile phase A consisted of 97:3 water/ methanol and mobile phase B was 100% methanol; both A and B contained tributylamine and glacial acetic acid at concentrations of 10mM and 15mM, respectively. The column was back-flushed with mobile phase C (100% acetonitrile, no additives) between injections for column cleaning.

The LC gradient was as follows: 0-2.5 min, 0% B; 2.5-7.5 min, linear ramp to 20% B min, 7.5-13 min, linear ramp to 45% B; 13-21 min linear ramp to 99% B and held at 99% B until 25 min. At 25 minutes, the 10-port valve was switched to reverse flow (back-flush) through the column, and the solvent composition changed to 95% C and held there for 3 min. From 28 to 28.5 min, the flow rate was ramped to 0.8 mL/min, held until 32.5 min, then reduced to 0.6mL/min. From 32.5 to 33.25 the solvent was ramped from 99% to 0% C while flow was simultaneously ramped down from 0.6-0.4mL/min and held until 39.9 min., at which point flow rate was returned to starting conditions at 0.25mL/min. The 10-port valve was returned to restore forward flow through the column at 40 min. An isocratic pump was used to introduce reference mass solution through the reference nebulizer for dynamic mass correction. Total run time was 30 min. The injection volume was 5 uL.

Data was analyzed using Mass Hunter Profinder 8.0 and Skyline. For all the cell lines, tissue and plasma unlabeled samples were used as controls for accurate detection of methionine isotopologs.

Positive method: Samples were analyzed by hydrophilic interaction liquid chromatography (HILIC)-MS in positive ion mode using an adapted version of a previously published method.^45^ Dried samples for HILIC analysis were reconstituted in 85/15 acetonitrile water. LC-MS runs were performed using a Thermo Vanquish UHPLC coupled to an Orbitrap ID-X mass spectrometer. The chromatographic column was a Waters BEH Amide column (2.1 x 100 mm ID, 1.7 µm). Mobile phase A and B were composed of 95/5 and 5/95 water/acetonitrile, respectively; both were modified by addition of 10mM ammonium formate and 0.125% formic acid. For mobile phase B, the ammonium formate must first be dissolved in the aqueous portion, followed by addition of acetonitrile and sonication for 10 minutes until the solution becomes clear. The column flow rate was 0.3 mL/min, the column temperature was 55° C and the injection volume was 5 µL. The gradient was as follows: 0-0.5min, 100%B, 0.5-7min, linear ramp 100-85%B, 7-9 min, 85%B, 9-16min, linear ramp 85-50%B, 16-16.1min, linear ramp 50-100%B, 16.1-20min, 100%B. MS source parameters were as follows: source type H- ESI, spray voltage +3200V, sheath gas 40, aux gas 10, ion transfer tube temperature 325° C, vaporizer temperature 300° C. MS data acquisition parameters were as follows: Data acquisition mode: MS1 scan (profile), orbitrap resolution 120k, scan range m/z 70-800, maximum injection time 50 ms, AGC target 100,000, Normalized AGC target 25%, microscans = 1, RF lens 45%, ETD internal calibration ON. Relative quantitation by peak area was performed using Skyline^46^, with peaks assigned based on accurate mass and retention time determined by analysis of authentic standards.

In Fig. 7D and Fig. S2G- S2I, a different column, the Intrada Amino Acid (Imtakt USA, Portland, OR) was used for HILIC chromatography in attempt to improve column lifespan and MS method details are in supplementary methods.

*Steady state metabolomics*: LC-MS analyses were performed using an Agilent Technologies Triple Quad 6470 LC-MS/MS system with a 1290 Infinity II LC Flexible Pump (Quaternary Pump), 1290 Infinity II Multisampler, 1290 Infinity II Multicolumn Thermostat with 6 port valve and 6470 triple quad mass spectrometer. Agilent MassHunter Workstation Software LC/MS Data Acquisition for 6400 Series Triple Quadrupole MS with Version B.08.02 was used for compound optimization and sample data acquisition. Agilent MassHunter Quantitative Analysis B.09.00 QqQ software was used to integrate and quantitate areas.

There are two identical Agielnt 1290 UHPLC systems-LC1 just for ion pairing chromograph method-neg mode MS/MS and LC2 is for other chromograph methods in both positive and negative mode MS/MS methods.

In LC1, 220 metabolites are from Agilent ion pairing method dedicated to the LC1-6470 QqQ MS. The LC-MS/MS method was created with dynamic MRM (dMRM) with RTs, RT windows from authentic standards in Agilent application lab. In addition, we add 10 more compounds into the existing methods in negative MS with the same chromograph method with authentic compounds.

In LC2, retention time (RT) and dMRMs for each metabolite was measured from a pure standard solution or a mix standard solution for all positive and/or neg MS/MS mode methods.

In LC2, positive acquisition mode, a Waters Acquity UPLC BEH TSS C18 column (2.1 × 100mm, 1.7µm) with ACQUILITY UPLC C181.7µm VanGurad Pre-Colum was used with mobile phase (A) consisting of 0.1% formic acid in water; mobile phase (B) consisting of 0.1% formic acid in acetonitrile. Gradient program: mobile phase (B) was held at 1% for 3 min, increased to 6% in 12 min, then to 15% B in 15 min, then to 99% B in 17 min and held for 2 min before going to the initial condition at 19.1 min and held for 5 min. The column was at 40 C and 3 µl of sample was injected into the LC-MS with a flow rate of 0.2 ml/min. Calibration of TOF MS was achieved through Agilent ESI-Low Concentration Tuning Mix. Key parameters of electrospray ionization (ESI) are: Gas temp 275 C, Gas Flow 14 l/min, Nebulizer at 20 psi, SheathGasHeater 250 C, SheathGasFlow 11 l/min, Capillary 3000 V. MS: Delta EMV 200 V, Cycle Time 500 ms, Cell Acc 4 V.

In LC1, the negative acquisition mode, Agilent ZORBAX RRHD Extend-C18, 2.1 × 150 mm, 1.8 µm and ZORBAX Extend Fast Guards for UHPLC are used in the separation. Ion paring (IP) stock solution is prepared by mixing GC-grade 450 ml of methanol, 35.8 ml of Tributylamine (TBA) and 12.9 ml of acetic acid. Solution A is prepared with 1000 ml of HPLC grade water, 1034 ul of deactivator and 34.25 ml of IP stock solution. Solution B is prepared with 1000 ml of HPLC grade methanol, 1034 ul of deactivator and 34.25 ml of IP stock solution. Solvent A is 97% water and 3% methanol with 15 mM acetic acid and 10 mM TBA at pH of 5. Solvent C is 15 mM acetic acid and 10 mM TBA in methanol. Washing Solvent D is acetonitrile. LC system seal washing solvent is 90% water and 10% isopropanol. Needle washing solvent is 50% methanol, 50% water. LC gradient profile is: at 0.25 ml/min, 0-2.5 min, 100% A; 7.5 min, 80% A and 20% C; 13 min 55% A and 45% C; 20 min, 1% A and 99% C; 24 min, 1% A and 99% C; 24.05 min, 1% A and 99% D; 27 min, 1% A and 99% D; at 0.8 ml/min, 27.5-31.35 min, 1% A and 99% D; at 0.6 ml/min, 31.50 min, 1% A and 99% D; at 0.4 ml/min, 32.25-39.9 min, 100% A; at 0.25 ml/min, 40 min, 100% A. Column temp is kept at 35 C, samples are at 4 C, injection volume is 2 µl. 6470 Triple Quad MS is calibrated with ESI-L Low concentration Tuning mix. Source parameters are: Gas temp 150 C, Gas flow 10 l/min, Nebulizer 45 psi, Sheath gas temp 325 C, Sheath gas flow 12 l/min, Capillary - 2000 V, Delta EMV -200 V. Dynamic MRM scan type is used with 0.07 min peak width, acquisition time is 24 min. Delta retention time of plus and minus 1 min, fragmentor of 40 eV and cell accelerator of 5 eV are incorporated in the method. Data was analyzed using MassHunter QQQ quantitative analysis

### 13C5 methionine infusions

Tumor-bearing mice were surgically implanted with catheters in the jugular vein and carotid artery. After surgical recovery, the cannulated mice were starved for 5 hours and then anesthetized using 2% isoflurane inhalation and given cranial RT (Lead shield leaving the cranium exposed) or sham RT using an orthovoltage irradiator. Immediately after RT, bright, alert, and active mice were administered with a bolus (0.0135 mg/g bolus) followed by continuous infusion (0.00053 mg/g/min) for 60 mins. The jugular vein catheter was used for ^13^C_5_ methionine infusion, and the carotid artery catheter was used to collect plasma during infusions. The plasma was collected at serial time points in EDTA-coated tubes and flash-frozen immediately. At the end of the infusion, mice were given ketamine (50mg/Kg) intravenously to induce rapid anesthesia. Following euthanasia, the tissue samples were harvested as quickly as possible using fluorescence guide microdissection (to separate the GFP-expressing tumor tissue from the normal brain). All tissue samples were immediately flash frozen using liquid nitrogen for further analysis.

### Correction of mice inter-variability for U^13^C-methionine saturation

Since mice were euthanized at different time points, the enrichment of M+5 methionine in plasma might not have reached saturation at earlier time points. In addition, the saturation level might differ from mouse to mouse. To remove this inter-variability of plasma methionine enrichment between mice, we fitted the plasma methionine M+5 enrichment to an exponential decay function (Eq. 1) based on the tracer kinetics model described in ^47^. We observed that plasma methionine M+5 enrichment at 30 min reaches steady state and we chose this timepoint as a reference to estimate function parameters.

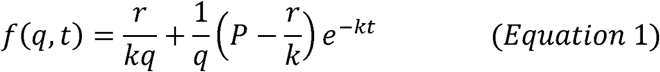

where the rate of tracer infusion, *r = 0.00053 mg/min/g* and the bolus infusion, *P = 0.0135 mg/min/g*. *t* is the time of infusion, *k* is the rate constant for methionine enrichment and *q* is the methionine pool size and may vary mouse to mouse. *k* and *q* are determined by fitting the experimental data. These fitted parameters are then used in Eq. 2 to calculate the saturation enrichment (*SE*) for each mouse.

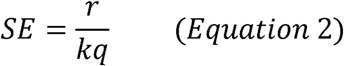

To remove the inter-variability in saturation enrichment of mice, a correction factor is calculated for each mouse by dividing the mouse saturation enrichment to the average of saturation enrichment values across mice. Then the labeled MIDs are divided by the correction factor.

### Metabolic model of the methionine pathway

Methionine model was implemented based on the active reactions occur in mouse from KEGG database^47^ and the availability of experimental enrichment data. Methionine model consists of six unidirectional reactions (Table 1) and three balanced metabolites (Table 2). Our experimental data has shown that cystathionine enrichment is low, but some labeling can be observed. Hence, we assumed that there is a net flux going out of methionine cycle to produce cystathionine. Since the enrichment data for homocysteine was not available, it was skipped in the cycle, but to account for the cystathionine production from homocysteine, a sink flux was added for SAH. Since MTAP is deleted in our experiments which catalyzes the production of methionine from MTA, we did not include this reaction in our model. However, since we observed labeling in MTA, we added a sink flux for SAM to account for a loss flux from the methionine cycle to produce MTA. Plasma methionine labeling was assumed to be M+5 only and used to normalize MIDs of tissue metabolites including methionine M+5 and M+4, SAM M+5, and SAH M+4. It is assumed that the methyl donor 3- methyltetrahydrofolate (MTHF) in SAH ➔ Met reaction is unlabeled.

**Table 1.** List of methionine model reactions. . Irreversible reaction refers to a reaction that proceeds in only one direction and sink refers to a reaction that consumes the metabolite outside of the model.

| Reaction No. | Reaction | Reaction Type |
| --- | --- | --- |
| $v_1$ | Plasma Met → Met | Irreversible |
| $v_2$ | Met → SAM | Irreversible |
| $v_3$ | SAH + MTHF → Met + THF | Irreversible |
| $v_4$ | SAM → SAH + CH<sub>3</sub> | Irreversible |
| $v_5$ | SAM → out | Sink |
| $v_6$ | SAH → out | Sink |

**Table 2.** List of metabolites and their experimental concentrations used in methionine model.

| Metabolite | Concentration in brain (mean ± standard deviation, nmol/g tissue) | Concentration in GBM (mean ± standard deviation, nmol/g tissue) |
| --- | --- | --- |
| Met | 1390 ± 100 (n = 7)<sup>52</sup> | 1930.41 ± 1002.34 |
| SAM | 13.0 ± 0.50 (n = 23)<sup>53</sup> | 16.42 ± 7.60 |
| SAH | 1.70 ± 0.16 (n = 23)<sup>53</sup> | 2.77 ± 1.42 |

### Estimation of metabolic fluxes in methionine model

We have previously adopted metabolic flux analysis^48,49^ for *in vitro* enrichment data and isotopic nonsteady state metabolic flux analysis (INST-MFA) to estimate metabolic fluxes of purine and pyrimidine reactions from *in vivo* enrichment data^50^. Here, a similar approach was used to estimate metabolic fluxes, concentration of balanced metabolites, and MIDs of balanced metabolites in methionine model. INST-MFA parameters including metabolic fluxes (*V*) and concentrations (*C*) are assumed to be at steady state. These parameters are generated randomly with 100 initial values and adjusted such that the objective function shown in Eq. 3 is minimized. Mean and standard deviation (*SD*) of metabolite concentrations in mouse brain are extracted from literature and corrected for GBM samples based on the ratio of total ion abundances in GBM samples over normal brain samples and the standard deviation is calculated by error propagation (Table 2). The objective function consists of two terms: (1) the squared standard score for estimated concentrations which tries to minimize the difference between experimental (*C_exp_*) and estimated concentrations (*C_est_*); (2) the squared standard score for estimated MIDs which tries to minimize the difference between experimental (*MID_exp_*) and estimated MIDs (*MID_est_*).

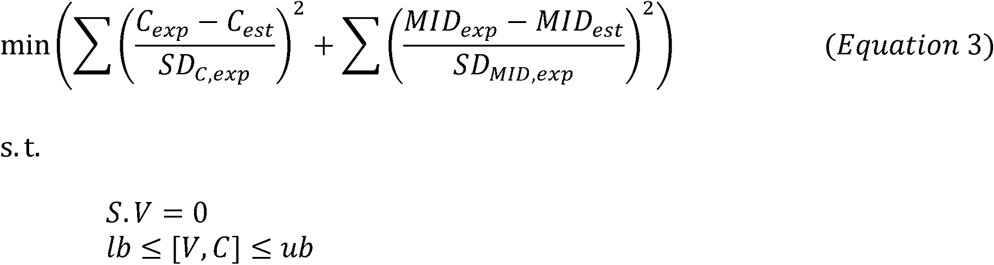

For each set of randomly generated parameters [*V,C*], a system of ordinary differential equations (time-dependent isotopologue balance equations) is solved to estimate MIDs of balanced metabolites (Eq. 4). MATLAB solver ode23s is used to solve for time-dependent MIDs as an initial value problem with the initial guess of experimental MIDs at 15min.

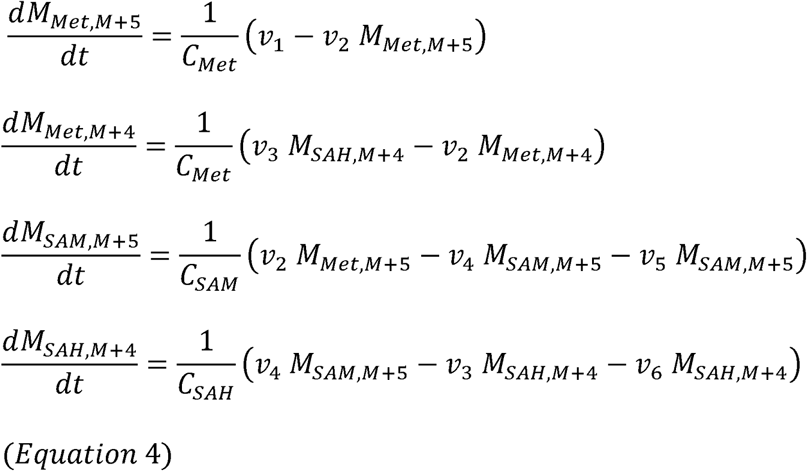

The optimization problem shown in Eq. 3 is also constrained linearly (*S.V* = 0) to ensure that estimated fluxes meet stoichiometry criteria (Eq. 5).

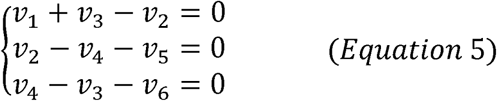

Finally, the optimization search space is defined within a lower and upper bound for each parameter. For fluxes, lower and upper bounds are set to [1, 500] for brain samples. The upper bound of fluxes is adjusted to 1000 so the optimal solution is within the optimization space and not on the boundary. The initial guesses of fluxes are also adjusted to ensure that the solution is not overfitted. At convergence, the minimized objective function value (Eq. 3) is a stochastic variable with a *χ*^2^-distribution. With 95% confidence level for the goodness of fit, the objective value is adequate if it falls in *χ*^2^(*n*,2.5%),*χ*^2^(*n*,97.5%)J where n is the degree of freedom.^51^ Solutions that meet this criterion are selected as feasible solutions and their mean and standard deviation were used in visualization of fluxes. Lower and upper bounds of concentrations are set to [*C_exp_* -*SD_c,exp_C_exp_* + *SD_c,exp_*] (Table 2). The optimization problem is solved using Artelys Knitro software (v12.4) in MATLAB R2021b.

### Clonogenic survival assay

Immortalized cells were either untreated or treated and irradiated with doses as indicated, followed by replating at clonal density. After 10-14 days, plates were fixed, and stained, and colonies (A colony is a group of 50+ cells) were counted. The enhancement ratio is calculated as the ratio of DMID (mean inactivation dose) of control and treatment. DMID refers to the linear range in a linear-quadratic survival curve.^54^

### Long term neurosphere formation assay

HF2303 primary neurospheres were treated with AGI-41998 or grown in media lacking methionine overnight followed by RT. The spheres were dissociated into single cells 24 hours post-RT and plated as single cells into 96 wells at a density of 2000 cells per well. After 7-10 days, the cell viability was assessed using CellTiter-Glo 3D reagent (Cat# G9682, Promega). Enhancement ratio is calculated as ratio of GI50 of control by treatment. GI50 is half maximal growth inhibitory concentration.

### Immunofluorescence

Immortalized or primary GBM cells treated and irradiated as indicated and fixed using 4% paraformaldehyde. The spheres were mixed with Histogel (Cat# HG-4000- 012, Thermo Scientific), transferred into tissue cassettes, and then embedded in paraffin for sectioning followed by staining. Mouse monoclonal antibody anti-phospho- Histone H2AX (1:1000 dilution; Cat # 9718S, Cell Signaling) and goat anti-mouse IgG Alexa fluor 594 secondary antibody, (1:2000 dilution; Cat# A-11005, Invitrogen) were used for γ-H2AX foci staining. A threshold of 10 or 5 foci per nucleus is counted as a positive cell for γ-H2AX for immortalized cells or neurospheres respectively.

### Animal Studies and xenograft models

All the animal experiments were approved by the University Committee on Use and Care of Animals at the University of Michigan. All the mice were housed in specific pathogen-free conditions. Animals were housed at 74°F temperature with a relative humidity range of 30-70% and a 12-hour light/dark cycle with unlimited access to food (PicoLab® Laboratory Rodent Diet, 5L0D) and water. All animals used in the experiments were 4-12 weeks old. RT-resistant GBM6 and GBM38 PDX models were obtained from Dr. Jann Sarkaria (Mayo Clinic National PDX resource). These PDX models were then propagated as subcutaneous flank tumors in female CB17-SCID mice. These tumors were subcutaneously implanted bilaterally into the dorsal flanks of CB17-SCID mice. Once the tumors reached 80-100 mm^3^ the mice were randomized into 4 groups, Control (1% w/w HPMCAS-CF, 1% w/w PVP K30, 2% w/w TGPS, 0.1% simethicone in ultrapure water), RT(2 Gyx5), AGI-41998(30 mg/Kg), or AGI-41998+RT were administered as shown in Fig. 5A. A subset of tumors were harvested 6 hours after last second of RT (2 Gy) or sham RT for LC-MS analysis. Tumor volume and body weight were measured three times a week. Tumor volumes were determined using digital calipers and calculated using the formula (π/6) × Length × Width². End point tumors were harvested 6 hours after last dose of AGI-41998 or vehicle and analyzed by LC-MS.

### Orthotopic PDX models

For intracranial models, HF2303 or GBM6 (GBM6-after a brief period of explant culture) cells were infected with lentiviruses harboring GFP and luciferase (lenti-LEGO- Ig2-fluc-IRES-GFP-VSVG) and GFP positive cells were enriched by flow cytometry. To generate intracranial tumors, Rag1 mice were anesthetized, and a small incision was made on the scalp. A burr hole was created at coordinates of 1 mm forward and 2 mm lateral from the bregma. Around 3 × 10^5^ (GBM38) - 2 x 10^6^ (HF2303) cells were injected at a rate of 1 µL/min. To assess the intracranial tumor growth, the mice were intraperitoneally injected with 150mg/Kg of D-luciferin and 10 min later imaged using IVIS™ Spectrum imaging system (PerkinElmer) while under anesthesia (2% isoflurane inhalation). Bioluminescent flux values were used to randomize the mice into 4 treatment groups, Control, RT, AGI-41998, or AGI-41998+RT. Bioluminescence flux values for mice euthanized midway through experiments were estimated using the latest measurement before euthanasia. AGI-41998 was administered daily via oral gavage and mice were treated with RT using an orthovoltage irradiator following the regimen as shown in Fig. 6A and 6F. In the GBM6 model, we have chosen to administered AGI-41998 at a dose of 15 mg/kg to mitigate the combined weight loss caused by both the tumor and the treatment.

### Statistical analysis

Statistical analysis was performed using GraphPad Prism. A students t-test was used to compare two groups. For compare three or more groups, a one-way ANOVA was used followed by multiple comparison tests such as Dunnett’s multiple comparisons test or Tukey’s multiple comparisons test appropriately. Statistical significance for survival analysis was performed Kaplan–Meier method and compared using Log-rank (Mantel-Cox) test.

## RESOURCE AVAILABILITY

### Lead contact

Daniel R Wahl

### Data and code availability

Code generated in this manuscript will be publicly available on GitHub upon acceptance for publication.

### Funding and acknowledgements

We thank Dr. Jann Sarkaria for providing patient-derived xenograft models used in animal studies. This study was supported by Agios/Servier pharmaceutical grant, Rogel Cancer grant, Ben and Catherine Ivy Foundation. Rogel Comprehensive Cancer Center (P30CA046592), University of Michigan Animal phenotyping core (DK020572, DK089503, 1U2CDK110768). A.J.S was supported by NCI F32CA260735, S. P was supported by PF-23-1077428-01-MM, M.A.M, T.S.L, and D.R.W were supported by the SPORE grant, P50CA269022. M.A.M was supported by R01CA240515, D.R.W. was supported by the NCI (K08CA234416 and R37CA258346), Cancer Center Support Grant P30CA46592, Damon Runyon Cancer Foundation, Ben and Catherine Ivy Foundation, and the Sontag Foundation. Some illustrations were created using BioRender software

### Disclosures

D.R.W has consulted for Agios Pharmaceuticals, Admare Pharmaceuticals, Bruker and Innocrin Pharmaceuticals. DRW is an inventor on patents pertaining to the treatment of patients with brain tumors (U.S. Provisional Patent Application 63/416,146, U.S. Provisional Patent Application 62/744,342, U.S. Provisional Patent Applicant 62/724,337). AJS and CAL are co-inventors on U.S. Provisional Patent Application 63/416,146. In the past three years, C.A.L. has consulted for Astellas Pharmaceuticals, Odyssey Therapeutics, Third Rock Ventures, and T-Knife Therapeutics, and is an inventor on patents pertaining to Kras regulated metabolic pathways, redox control pathways in pancreatic cancer, and targeting the GOT1-ME1 pathway as a therapeutic approach (US Patent No: 2015126580-A1, 05/07/2015; US Patent No: 20190136238, 05/09/2019; International Patent No: WO2013177426-A2, 04/23/2015

