## Supplementary for "Reciprocal links between methionine metabolism, DNA repair and therapy resistance in glioblastoma"

### SUPPLEMENTARY METHODS:

#### Combination studies

For the combination studies, the U87 or KP4 cells were plated at a density of 2000 cells per well in a 96-well plate. After treatment with indicated agents, cell viability was assessed using cell titer glo assay. The combination index (CI) was calculated using the Chou-Talalay model.[1] The lowest CI was identified by treating the cells with multiple dose combinations of the two drugs. CI <1, indicates a synergistic effect, CI>1 indicates an antagonistic effect and CI=1 indicates an additive effect between any two given drugs.

#### Stable cell lines

Human MTAP, a gift from Vern Schramm (Cat # 64077, Addgene), was used as a template for PCR to generate the 6XHis-TEV-MTAP fragment used for Gibson assembly using Q5 High Fidelity DNA Polymerase (NEB). The MTAP PCR fragment was isolated by gel band isolation from a 1% TAE Agarose gel using the Monarch Gel Isolation Kit (NEB). The purified 6XHis-TEV-MTAP PCR product was cloned then into the BamHI/NotI sites of the lentiviral vector, pLentiLox RSV-CMV Puro, using Gibson Assembly. Clones were then sequenced to determine verify 6XHis-TEV-MTAP correct sequence and insertion into pLentiLox RSV CMV-Puro. The resulting clone was then named pLentiLox RSV 6xHis-TEV-MTAP -CMV Puro. This plasmid was then used for the generation of lentivirus using standard methodology. U87 and HF2303 were infected with lentiviral particles (LentiLOx-RSV-6HisTEV-MTAP-CMVpuro-VSVG) of human MTAP (Cat# 64077,

Add gene) and selected with puromycin at 1 µg/mL for 3 days. Pooled cells were harvested and MTAP expression was confirmed by western blotting.

**Mass Spectrometry analysis**

In Fig. 7D and Fig. S2G- S2I, the column dimensions were 10 cm x 3.0 mm inner diameter (Imtakt USA, Portland, OR). The column temperature was set at 37°C, the flow rate was 0.5 mL/min, and the injection volume was 10 µL. Mobile phase A was 100 mM ammonium formate in 80/20 water/acetonitrile and mobile phase B was 0.3% formic acid in acetonitrile. The chromatographic gradient was as follows: 0-4min, 80%B; 4min-14 min, linear ramp to 0%B; 14-16 min, 0%B, 16-16.1min, linear ramp to 80%B, 16.1-20min, 80%B. The instrumentation and all MS source parameters were identical to the previous HILIC method. MS data acquisition parameters were as follows: Data acquisition mode: MS1 scan (centroid), orbitrap resolution 60k, scan range m/z 100-800, maximum injection time 50 ms, AGC target 400,000, Normalized AGC target 100%, microscans = 1, RF lens 35%, ETD internal calibration ON. Relative quantitation and compound identification were performed identically to the previous HILIC method.

**Western Blotting**

Cells were lysed using RIPA lysis buffer (Cat# 89900, Thermo Scientific) supplemented with PhosSTOP phosphatase inhibitor (Cat# 04906845001, Roche) and complete protease inhibitor tablets (Cat# 1187358001, Roche). Proteins were detected using MTAP (Cat# 62765, Cell signaling), Vinculin (Cat# 13901S, Cell signaling), Histone H3 (Cat# 96C10, cell signaling).

**Tissue microarray**

Formalin-fixed, paraffin-embedded tissue blocks (FFPE) of primary brain tumor cases were obtained from the files of the Department of Pathology, University of Michigan Medical Center, Ann Arbor, MI. The University of Michigan Institutional Review Board provided a waiver of informed consent to obtain these samples (HUM00165469). The GBM patient blocks were reviewed by a clinical pathologist (SF), who marked the tumor area on each block, a tissue microarray was constructed from the most representative area using the methodology of Nocito et al.[2] Each case was represented by two 1 mm diameter cores, obtained from the most representative, non-necrotic area of the tumor. These were arrayed along with normal brain controls, other control tissue (liver, pancreas, etc) and colored orientation cores. Additionally, the TMA block is baked at least overnight at 37C, face down on a slide. Afterword, I set it in the PCR machine programmed to cycle for 1 min at 37C, 10 min at 52C, and then hold at 4C for at least 5 min.

**Immunohistochemistry**

Immunohistochemistry (IHC) was conducted using the ABC Vectastain kit (Cat# PK-6101) following the manufacturer's instructions. After deparaffinization, rehydration, antigen retrieval, and blocking, the tissue microarray (TMA) slide was incubated overnight at 4°C with the MTAP primary antibody (1:250 dilution, Cat# 62765, Cell Signaling). Following this, the slide was incubated with a secondary rabbit antibody for 30 minutes, then stained using the DAB kit (Cat# SK-4100) and counterstained with hematoxylin. Finally, the slide was dehydrated and mounted.

**Patient data**

Metabolomic data from flash-frozen human glioblastoma tumor samples were sourced from our previously published work.[3] The samples were grouped according to the percentage of MTAP-positive cells for analysis, and one outlier in the MTA was excluded from the dataset.

**Orthotopic PDX models**

Methionase: PEGylated Methioninase was acquired from Dr. Everett Stone. The methioninase was administered at a dosage of 100mg/Kg via intraperitoneal injection every two days, starting one day before radiation therapy and continuing until the endpoint.

Methionine-restricted diet: The mice were randomized and given the control diet with 0.84% Methionine (Cat# A11051302Bi, Researchdiets) or methionine-restricted diet with 0.12% Methionine (A11051301Bi, Researchdiets) starting two-days prior to radiation therapy and continuing until the endpoint.

**Supplementary Figure legends:**

**Supp Fig. 1:** A). Untreated intracranial HF2303 xenografts and the normal mouse brain were harvested, and methionine metabolites were analyzed using LC/MS. Data represented as mean±SEM for N=4 biological replicates. P values were calculated in comparison with the control (Brain). B-D). HF2303 tumor bearing mice were treated with cranial RT(8GY) and tumor and brain were harvested 1 hour post-RT for LC-MS analysis.

E). GBM6 tumor bearing mice were treated with RT (8GY) and tissues were harvested 1 hour post-RT for MS analysis. P values were calculated in comparison with the control (Brain-RT). F). U87 or HF2303 flank tumors were irradiated (8GY) and harvested 2 hours post-RT for LC-MS analysis. A-F). **P ≤ 0.01 and ***P ≤ 0.001.

**Supp Fig. 2:** A). U87 cells were labeled with $^{13}C_5$ methionine and untreated or irradiated cells were harvested 1 hour post-RT for LC-MS analysis. Relative m+5 methionine levels in U87 cells from N=7 biological replicates. B-C). Relative m+4 methionine (recycled methionine levels) or relative m+4 SAM (recycled SAM) in either untreated or RT treated U87 cells from N=7 biological replicates. P value of remethylated SAM is 0.0334 D-F). HF2303 spheres were labeled with $^{13}C_5$ methionine and either untreated or radiated samples were harvested 1-hour post-RT for LC-MS analysis. Relative m+5 methionine, m+4 methionine (recycled methionine), m+4 SAM (recycled SAM) for N= 3-8 biological replicates. G-I). GBM6 spheres were labeled with $^{13}C_5$ methionine and either untreated or radiated (8GY) spheres were harvested 1-hour post-RT for LC-MS analysis. Relative m+5 methionine, m+5 SAM and m+4 SAH were plotted for N=5 biological replicates. J). Mice were infused with low, medium or high dose of $^{13}C_5$ methionine for 2.5 hours and relative arterial methionine levels were analyzed using LC-MS analysis. K-L). The labeling patterns of SAM and SAH in mouse brain after the 2.5 hours of $^{13}C_5$ methionine infusion were analyzed using LC-MS analysis. M**).** Absolute metabolic fluxes of recycled methionine in normal brain and GBM with or without RT from N=3 biological replicates. A-M). *P ≤ 0.05 and ****P ≤ 0.0001. N-Q). Estimated MIDs from isotopic nonsteady state methionine model using estimated fluxes and concentrations at optimal solution in normal brain and GBM with or without RT. The model minimizes the difference between experimental MIDs (orange dots) and estimated MIDs (blue lines) normalized by the standard deviation of experimental MIDs (orange error bars). The y-axis represents MIDs normalized by plasma methionine M+5.

**Supp Fig. 3:** A). Relative MTA levels in GBM tissue compared to the contralateral cortex in GBM38 and HF2303 GBM xenograft models. Relative MTA levels in GBM38 and HF2303 models were analyzed from N=6 and N=9 biological replicates respectively. B). Representative MTAP IHC images from the tissue microarray with 100%, <20% and 20-60% MTAP-positive cells. C). Relative MTA levels in human patients with 100%, <20% and 20-60% MTAP-positive cells. D-E). Western blot showing MTAP expression in U87 MTAP or HF2303 MTAP cells. F-I). Impact of combinatorial treatment of AG270 or AGI-41998 and Bleomycin on U87 and KP4 cells. Lowest combination index (CI) is indicated, CI<1 implies synergistic effect, CI>1 implies antagonistic effect and CI=1 indicates additive effect. A-C). *P ≤ 0.05 and ****P ≤ 0.0001.

**Supp Fig. 4:** A-D). Representative images of γ-H2AX foci staining in U87 cells (corresponding to Fig. 4A), HF2303 neurospheres (corresponding to Fig. 4B), U87 cells (corresponding to Fig.4C), and HF2303 neurospheres (corresponding to Fig. 4D). Cells or spheres were grown in media depleted of methionine or treated with AGI-41998, irradiated (4GY), fixed and stained for γ-H2AX foci at the time points indicated. N=3 biological replicates. Scale bar: 50 μm. E-H). Representative images of γ-H2AX foci staining in U87 MTAP cells (corresponding to Fig. 4I), HF2303 MTAP neurospheres (corresponding to Fig. 4J), U87 MTAP cells (corresponding to Fig. 4K), and HF2303 MTAP neurospheres (corresponding to Fig. 4L). Cells or spheres were grown in media depleted of methionine or treated with AGI-41998, irradiated (4GY), fixed and stained for γ-H2AX foci at the time points indicated.  N=3 biological replicates. Scale bar: 50 µm. I). U87 WT and MTAP cells were grown in media with methionine and depleted of methionine for 24 hours and analysed using western blot. J). U87 cells were either untreated of treated with AGI-41998 and/or supplemented with 500 µM SAM for 24hours and harvested. Proteins were separated by gel electrophoresis and blotted using indicated antibodies L). Representative images of U87 cells (corresponding to Fig. 4M) untreated or treated with AGI-41998 and/or supplemented with 500 µM SAM, irradiated (4GY), fixed and stained for γ-H2AX foci at the time points indicated. N=3 biological replicates. Scale bar: 50 µm.

**Supp Fig. 5:** A-B). Mice were treated with 30 mg/Kg or 60 mg/Kg AGI-41998 and plasma, flank tumor and brain were harvested 6 hours post AGI-41998 treatment for LC-MS analysis. P values were calculated in comparison with the 30 mg/Kg for AGI-41998 and vehicle for SAM respectively. C-D). Body weights of GBM6 and GBM38 flank tumor bearing mice. Data is presented as mean ± SEM for N=7-8 mice per group. E-F). End point GBM6 and GBM38 tumors were harvested 6 hours after last dose of AGI-41998 and analyzed by LC-MS. P values were calculated in comparison with the control (Vehicle). A-F). *P ≤ 0.05, **P ≤ 0.01 and ****P ≤ 0.0001.

**Supp Fig. 6:** A-B). HF2303 tumor bearing mice were randomized into 4 groups. Mice were given AGI-41998 (30 mg/Kg) oral gavage and one dose of RT (8GY) and harvested 1 hour after RT. Data is represented as mean ± SEM from 15 (Control, RT and AGI-41998+RT) to four (AGI-41998) data points. P values were calculated in comparison with the control. C-D). Kaplan-Meier survival curves of HF2303 and GBM6 tumor bearing mice. E-F). Body weights of HF2303 and GBM6 orthotopic tumor bearing mice. Data is presented as mean ± SEM for N=7-8 mice per group. Data is presented as mean ± SEM for N=6-8 (HF2303) and N=8 (GBM6) mice per group. G). Schematic of treatment schedule in GBM6 tumor bearing mice. GFP and luciferase expressing GBM6 cells were intracranially injected into mice and once detected by bioluminescent imaging, the mice were randomized into 4 groups, Control, Methioninase (100mg/Kg), RT (2GY*5) and methionase+RT. Methioninase was dosed at 100mg/Kg via intraperitoneal injection every 48 hours beginning one day before RT. RT was administered as 5 fractions of 2GY. H). Total flux values were averaged for each group and plotted as fold change with respect to day1 of treatment. Data is presented as mean ± SEM for N=8-9 mice per group. I). Kaplan-Meier curve of GBM6 tumor bearing mice**.** P values of RT vs methioninase+RT is 0.045. P values were calculated using Log-rank (Mantel-Cox) test. J). Mouse weight in each treatment group of GBM6 tumor bearing mice. Data is presented as mean ± SEM for N=8-9 mice per group. K-L). End point tumors from Fig. S6G were analyzed using LC-MS and methionine and SAM levels in Plasma, brain and tumor are plotted. Data is presented as mean ± SEM for N=3-6 samples per condition. P values were calculated in comparison with the control. M). Kaplan Meier curves of GBM38 tumor bearing mice were given control diet or methionine restricted diet and/or treated with RT. N). End point tumors from Fig. S6M were analysed using MS analysis. Data is presented as mean ± SEM for N=2-5 mice per group. P values were calculated in comparison with the control. the control diet. A-N). *$P \leq 0.05$, ***$P \leq 0.001$ and ****$P \leq 0.0001$.
