## Supplementary Figures for "Reciprocal links between methionine metabolism, DNA repair and therapy resistance in glioblastoma"

**Figure. 1: Supplementary : RT acutely alters GBM methionine metabolism**

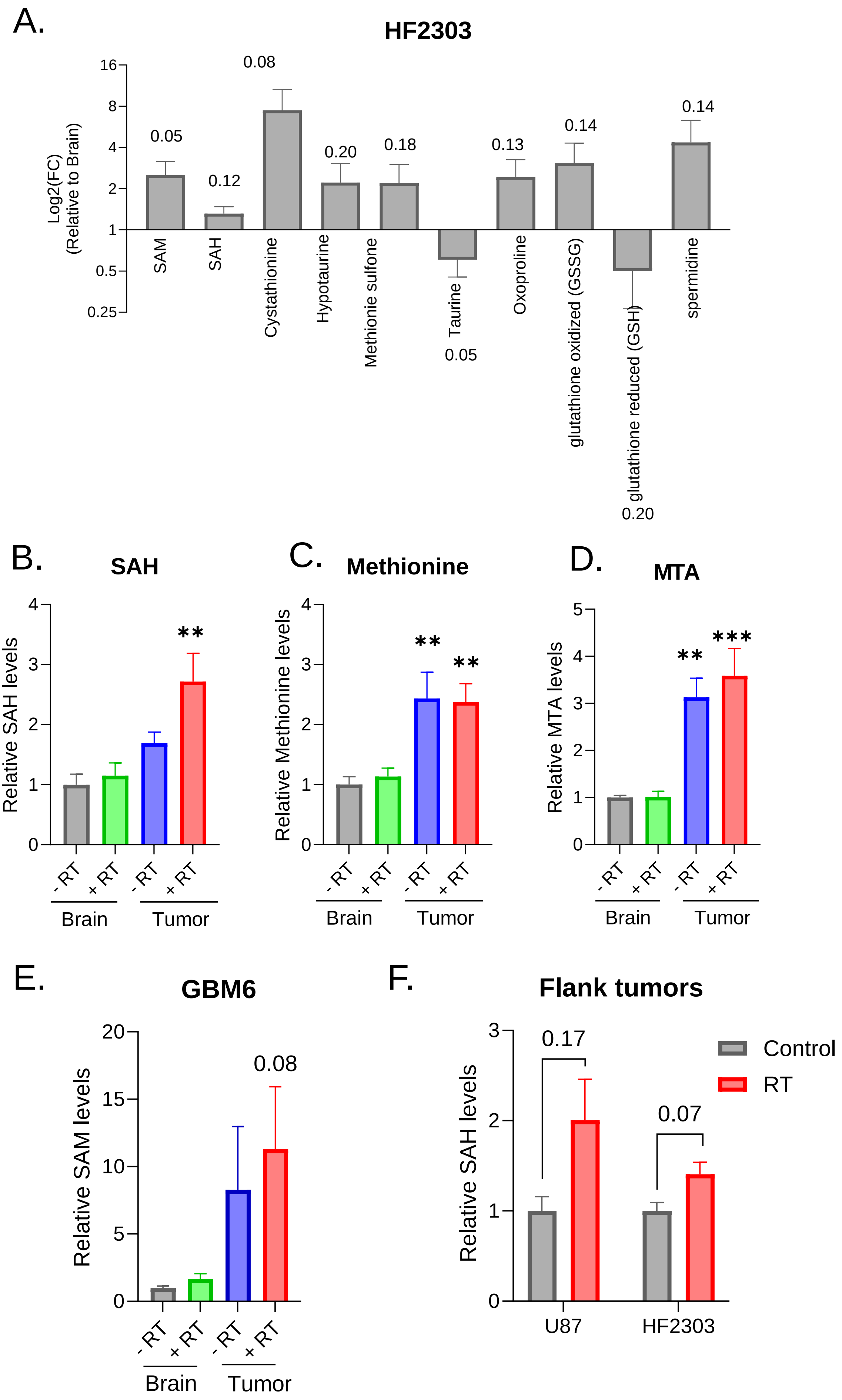

**Figure 2: Supplementary:RT acutely increases SAM synthesis in GBM cells and in intracranial GBM models**

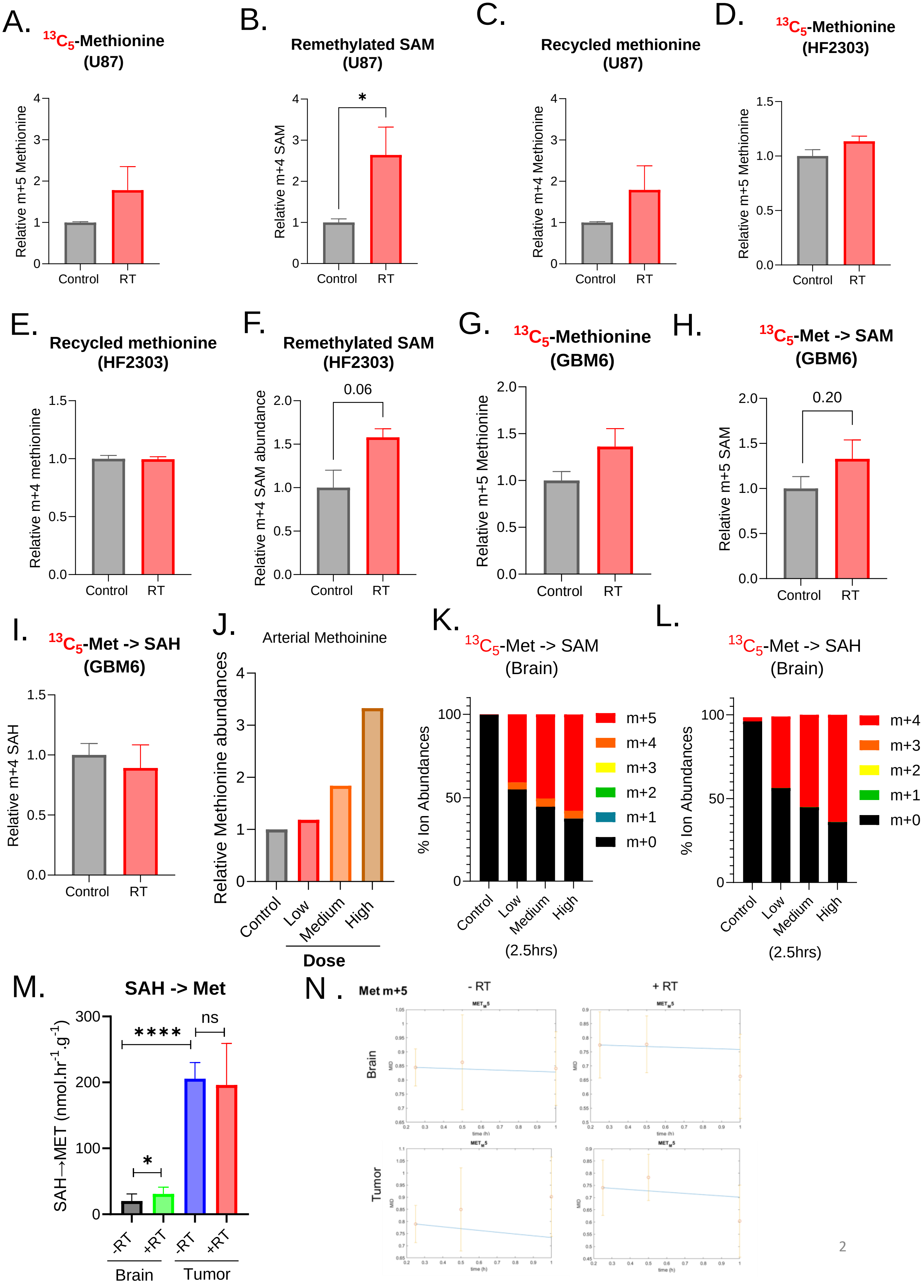

**Figure 2: Supplementary:RT acutely increases SAM synthesis in GBM cells and in intracranial GBM models**

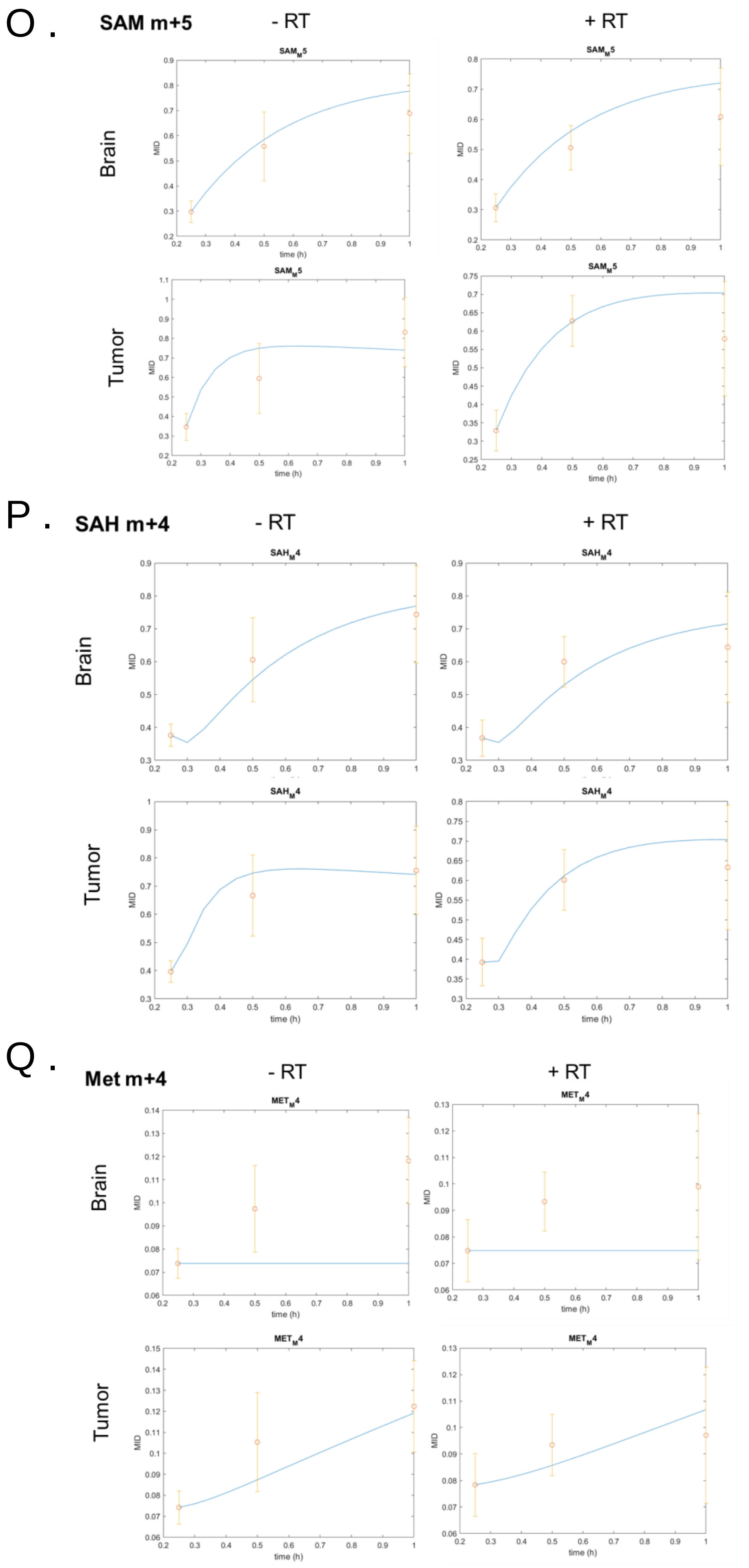

### Figure 3: Supplementary: Disrupting methionine metabolism enhances the responsiveness of GBMs to RT

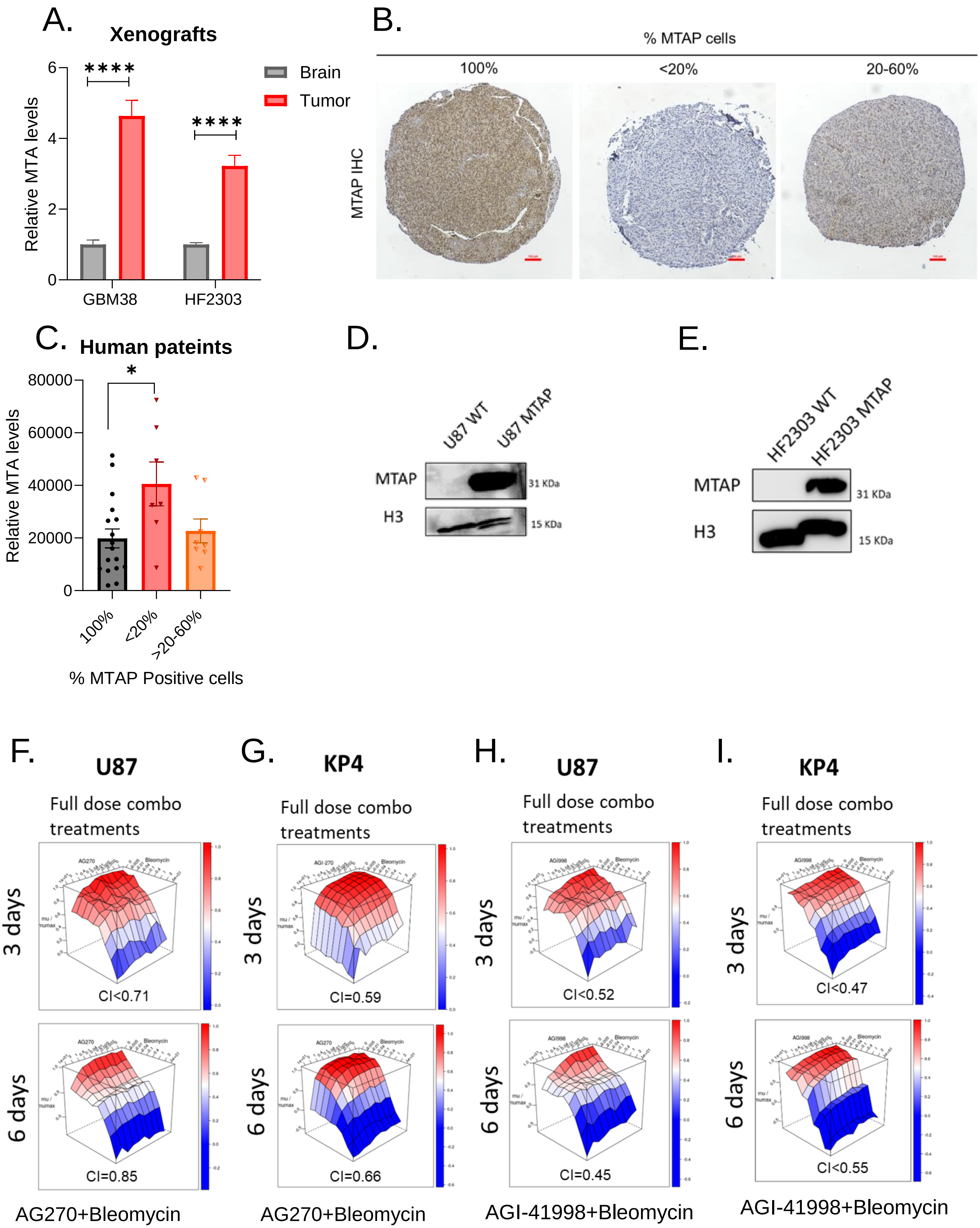

**Figure 4: Supplementary :Disrupting methionine metabolism slows DNA repair in GBM**

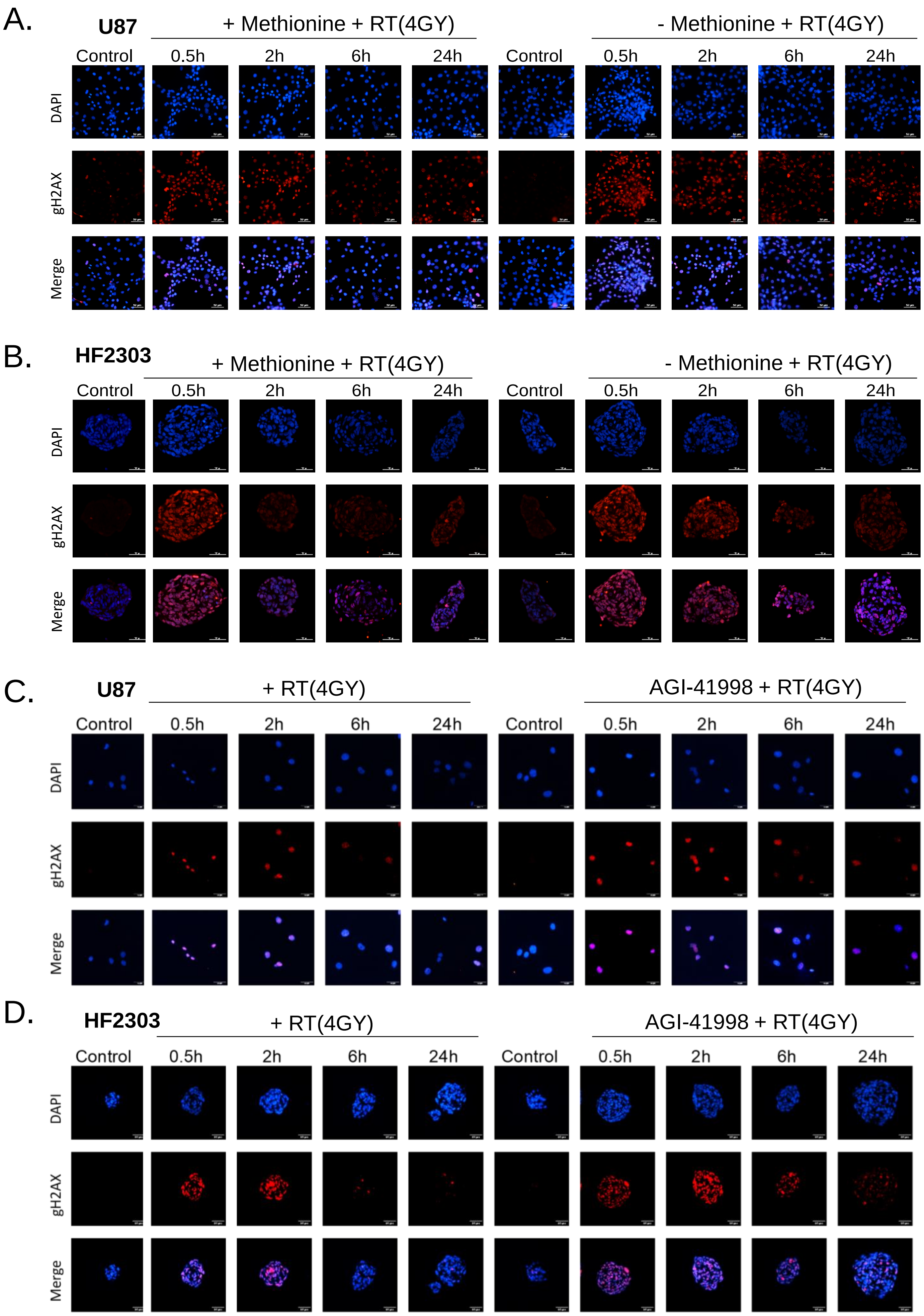

### Figure 4: Supplementary :Disrupting methionine metabolism slows DNA repair in GBM

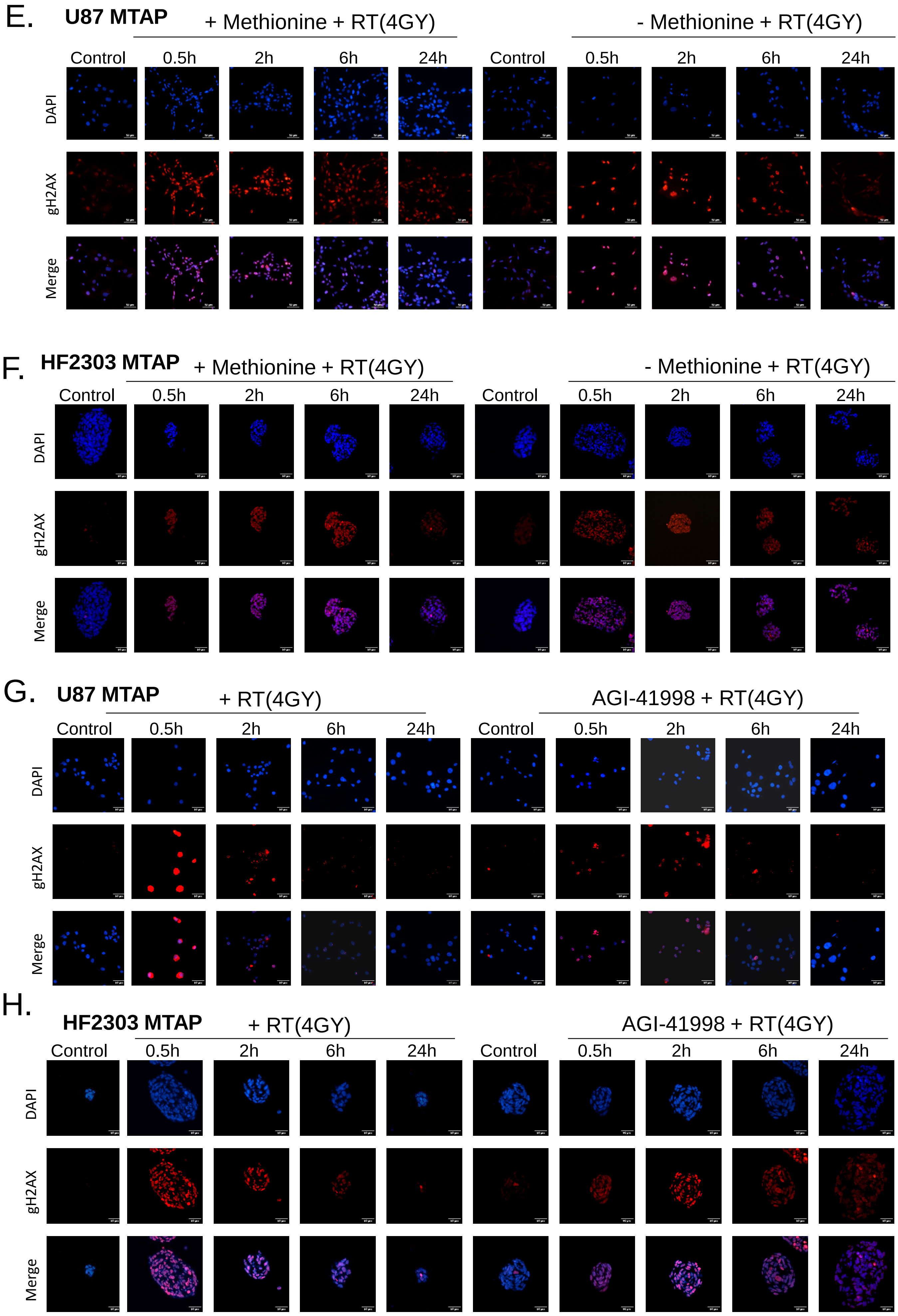

**Figure 4: Supplementary :Disrupting methionine metabolism slows DNA repair in GBM**

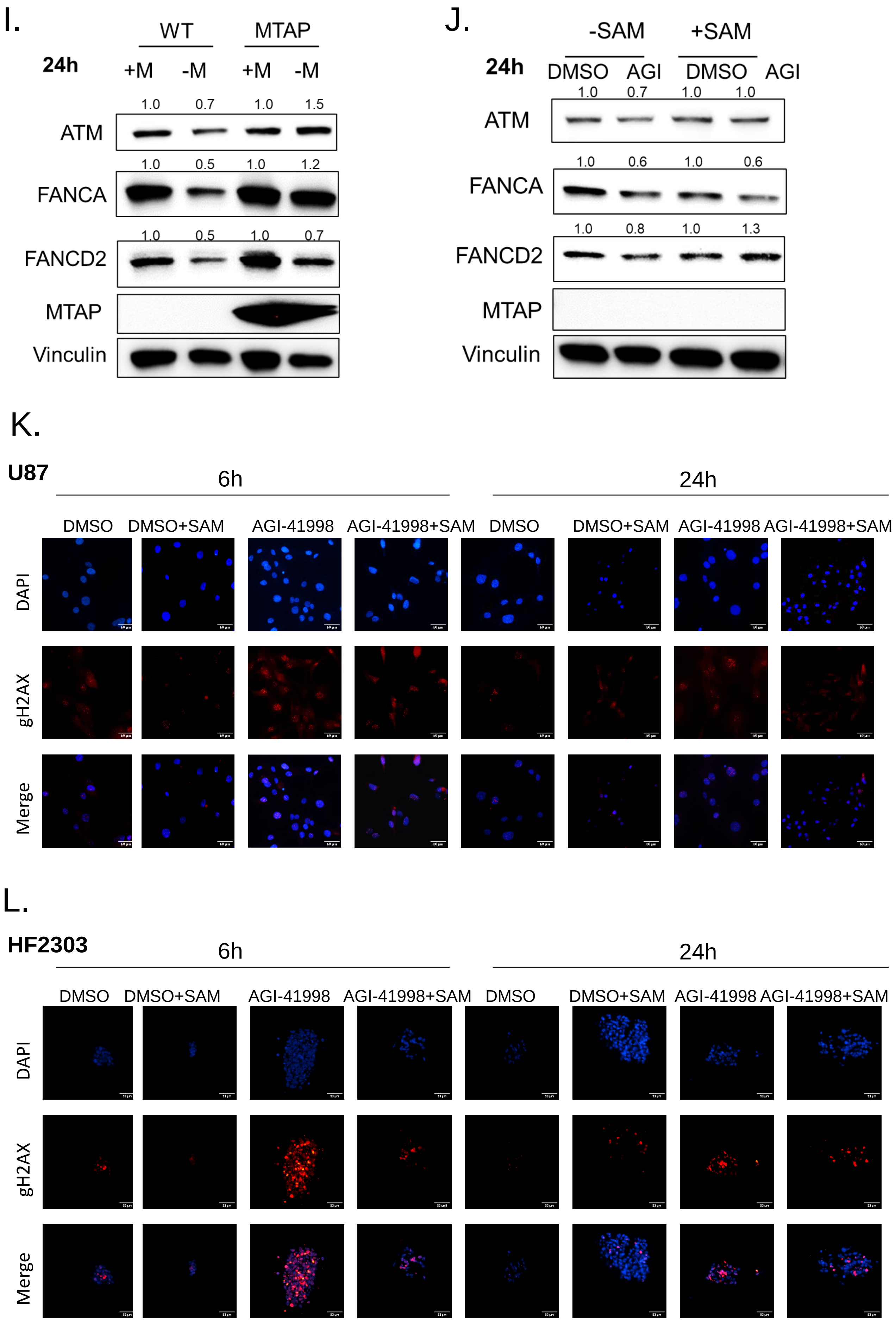

**Figure 5: Supplementary :MAT2A inhibition radiosensitizes MTAP-deleted flank models of GBM**

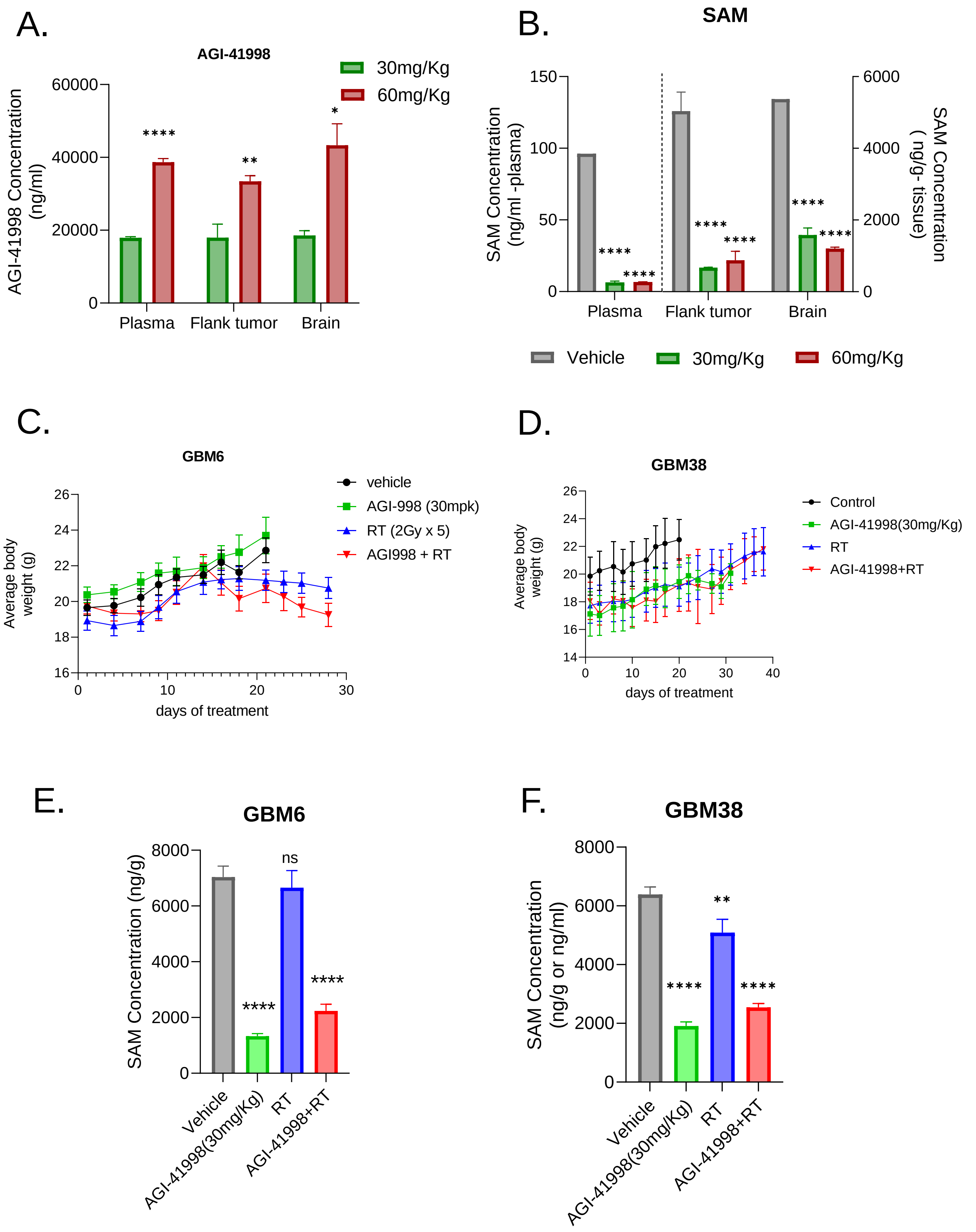

**Figure 6: Supplementary: MAT2A inhibition radiosensitizes intracranial models of GBM.**

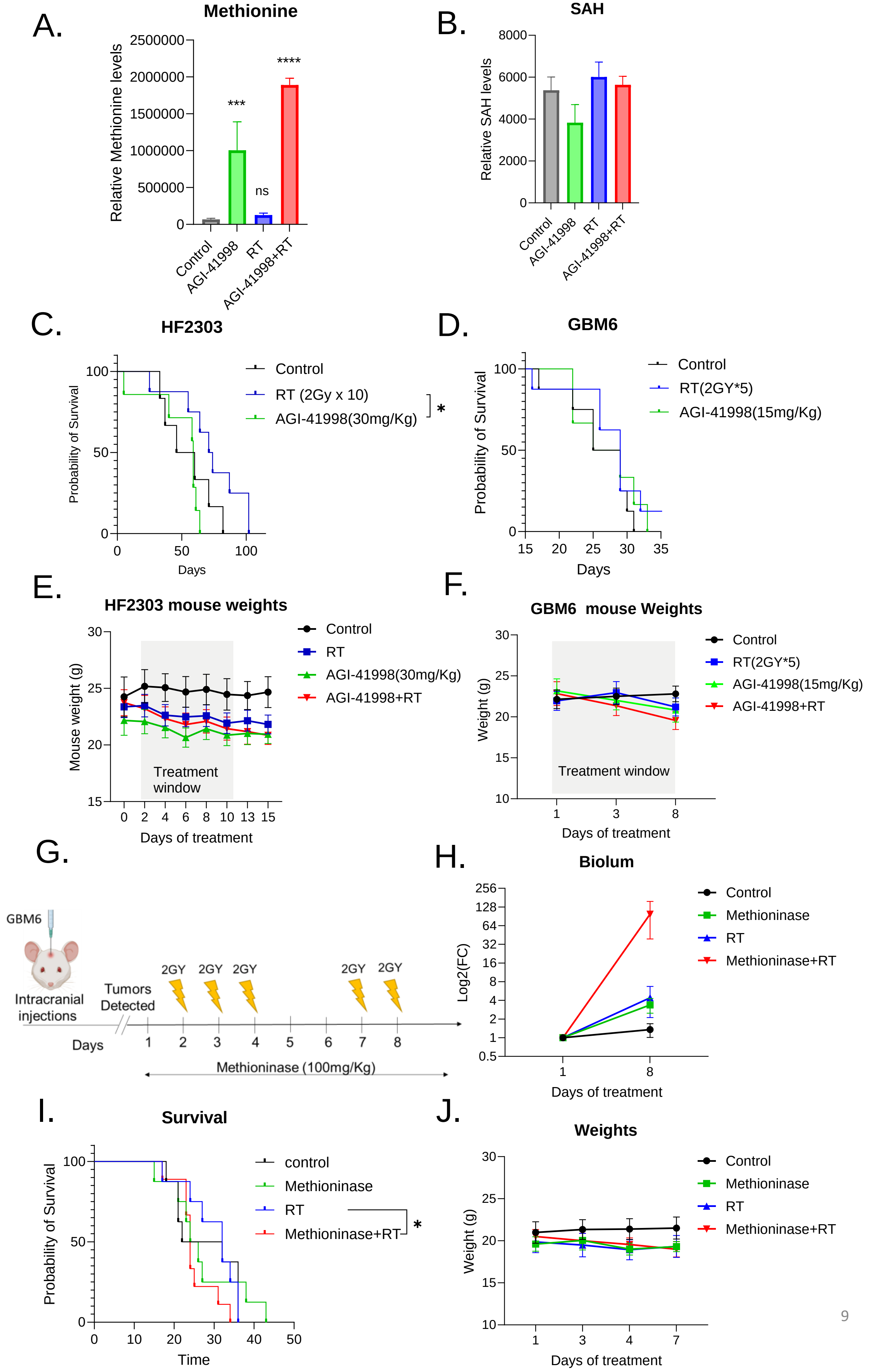

**Figure 6: Supplementary : MAT2A inhibition radiosensitizes intracranial models of GBM.**

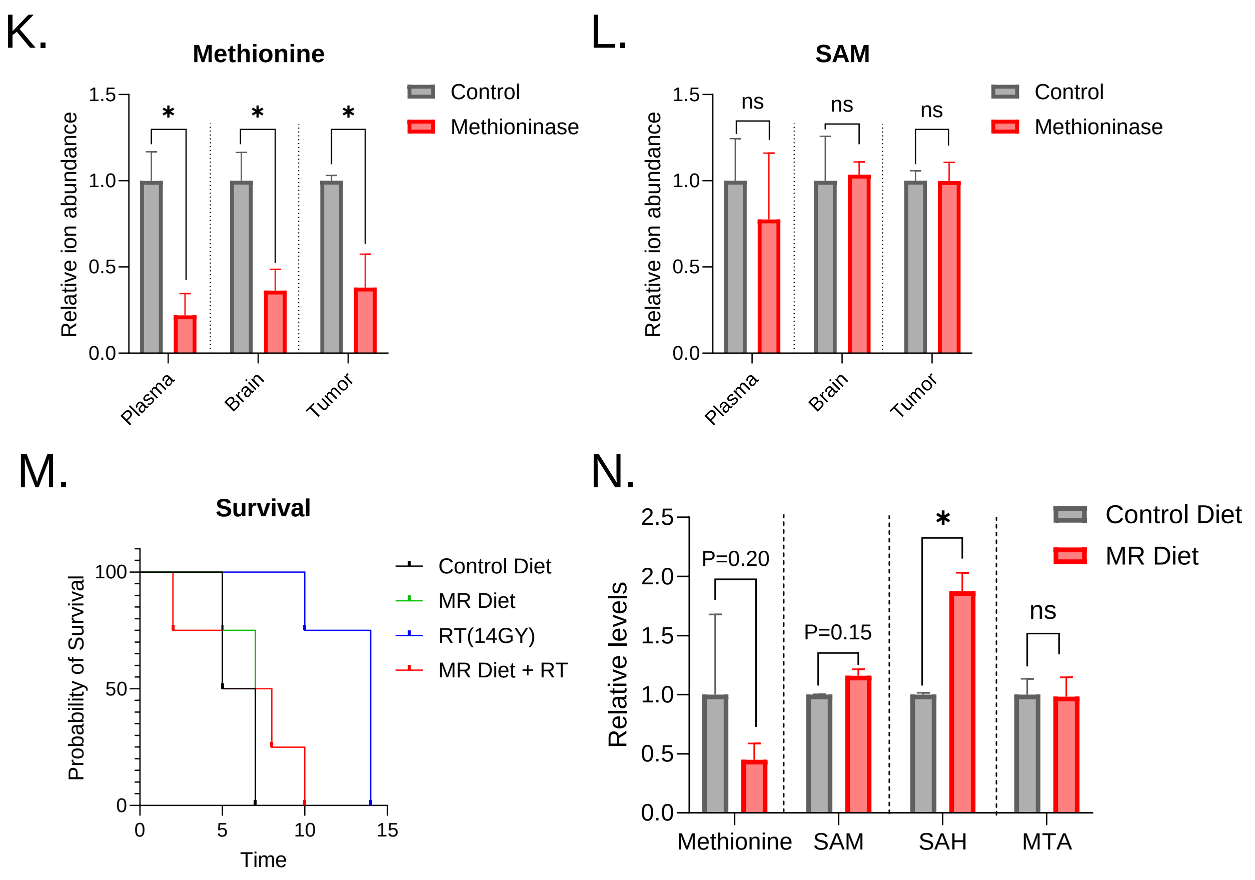
